# Functional dissection of the Pol V large subunit CTD in RNA-directed DNA methylation

**DOI:** 10.1101/111831

**Authors:** Jered M. Wendte, Jeremy R. Haag, Jasleen Singh, Anastasia McKinlay, Olga M. Pontes, Craig S. Pikaard

## Abstract

Plant multisubunit RNA Polymerase V transcription recruits Argonaute siRNA complexes that specify sites of RNA-directed DNA methylation (RdDM) for gene silencing. Pol V's largest subunit, NRPE1, evolved from the largest subunit of Pol II but has a distinctive carboxyl-terminal domain (CTD). We show that the Pol V CTD is dispensable for catalytic activity *in vitro*, yet essential *in vivo.* One CTD subdomain (DeCL), is required for Pol V function at virtually all loci. other CTD subdomains have locusspecific effects. In a yeast two-hybrid screen, the 3'->5' exoribonuclease, RRP6L1 was identified as an interactor with the DeCL subdomain and DeCL and glutamine-serine-rich (QS) subdomains, located downstream from an Argonaute-binding repeat subdomain. Experimental evidence indicates that RRP6L1 trims the 3’ ends of Pol V transcripts sliced by ARGONAUTE 4 (AGO4), suggesting a model whereby the CTD enables the spatial and temporal coordination of AGO4 and RRP6L1 RNA processing activities.

## Introduction

NUCLEAR RNA POLYMERASE V (Pol V) is a plant-specific multisubunit RNA polymerase important for siRNA-directed DNA methylation (RdDM) and transcriptional gene silencing, primarily of transposable elements (reviewed in (Haag and Pikaard 2011; Ream et al. 2013; Zhou and Law 2015). In the major RdDM pathway (reviewed in (Matzke and Mosher 2014; Wendte and Pikaard 2016), NUCLEAR RNA POLYMERASE IV (Pol IV) and RNA-DEPENDENT RNA POLYMERASE 2 (RDR2) generate short double-stranded RNAs that are then diced into 24 nt short interfering RNAs (siRNAs) (Blevins et al. 2015; Li et al. 2015; Zhai et al. 2015). These siRNAs associate with an Argonaute family protein, primarily AGO4 or AGO6, and guide cytosine methylation at sites of Pol V transcription. Pol V also has functions independent of 24 nt siRNAs (Pontes et al. 2009), including a pathway for establishing DNA methylation using 21 nt siRNAs derived from the degraded mRNAs of active transposons (Nuthikattu et al. 2013; McCue et al. 2015).

Pol IV and Pol V in Arabidopsis and maize are composed of twelve subunits, like Pol II (Ream et al. 2009; Ream et al. 2013; Haag et al. 2014). Approximately half of the Pol IV and Pol V subunits are encoded by the same genes as Pol II subunits (Ream et al. 2009; Ream et al. 2013). Remaining subunits, including the catalytic subunits (the two largest subunits), are encoded by genes that arose through duplication and sub-functionalization of ancestral Pol II subunit genes (Luo and Hall 2007; Ream et al. 2009; Tucker et al. 2010; Huang et al. 2015; Wang and Ma 2015). Arabidopsis Pols IV and V have three different subunits, but the only fundamental difference between maize Pols IV and V is their distinctive largest subunits, NRPD1 or NRPE1, respectively (Haag et al. 2014). NRPD1 and NRPE1 differ primarily at their C-terminal ends. NRPD1 has a short C-terminal domain (CTD) of ~100 amino acids that shares sequence similarity with a chloroplast protein implicated in plastid ribosomal RNA processing, DEFECTIVE CHLOROPLASTS AND LEAVES, (Keddie et al. 1996; Bellaoui et al. 2003; Bellaoui and Gruissem 2004; Haag and Pikaard 2011; Huang et al. 2015). Because of this similarity, this domain is abbreviated as DeCL (see Figure 1A). The Pol V largest subunit, NRPE1 has a much longer CTD, spanning ~700 amino acids. At the very C-terminus is a subdomain rich in repeated glutamine-serine (QS) motifs. The QS subdomain is preceded by a DeCL subdomain, similar (but not identical) to the DeCL domain of Pol IV. N-terminal to the DeCL domain is a subdomain consisting of ten 17 amino-acid repeats rich in glycine-tryptophan (GW) or WG dipeptide "ago-hooks" that bind to Argonaute family proteins, including AGO4 and AGO6 (El-Shami et al. 2007; Pontier et al. 2012). A fourth subdomain, termed the linker, is located between the 17aa repeats and the conserved sequences (domains A-H) that are characteristic of all multisubunit RNA polymerase largest subunits (see Figure 1A).

**Figure 1.**
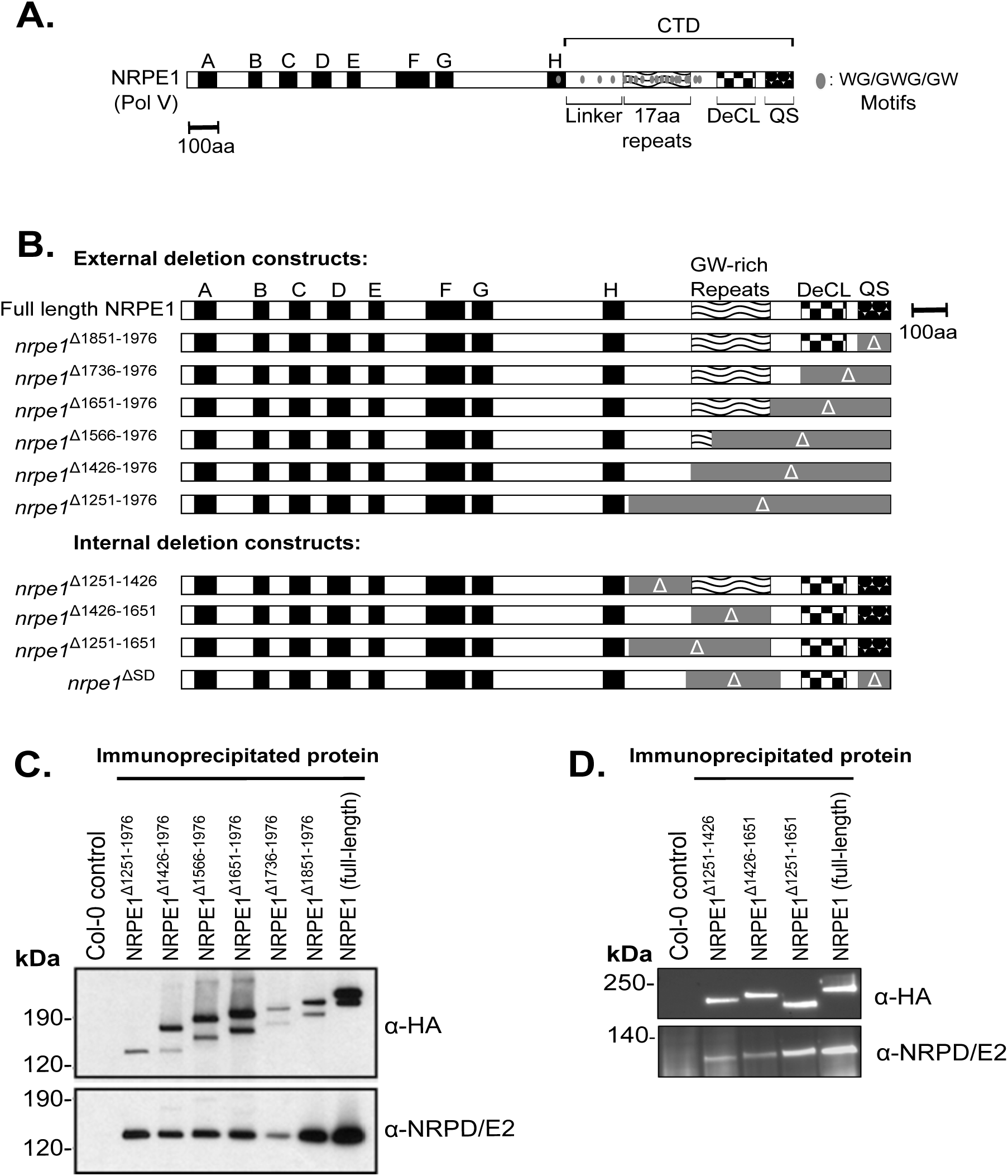
Engineering and expression of Pol V largest subunit (NRPE1) proteins with deletions in the C-terminal domain (CTD). **A.** Diagram of the NRPE1 protein of *Arabidopsis thaliana* (ecotype Col-0), showing the relative positions of domains A-H, conserved among multisubunit RNA polymerase largest subunits, and subdomains of the CTD. **B.** Diagrams of recombinant NRPE1 constructs tested in the study. All constructs were engineered to have a C-terminal HA epitope tag, except ΔSD, described previously (El-Shami et al. 2007). Numbering for deletion constructs denotes amino acid positions measured from the N-terminus. **C-D.** Immunoblot assays following anti-HA immunoprecipitation of C-terminal (**C**) or internal (**D**) CTD deletion constructs, with non-transgenic Col-0 serving as a control. Blots were probed using antibodies recognizing the HA epitope tag on recombinant NRPE 1 proteins, or recognizing the second-largest subunit used by both Pol IV and Pol V, NRP(D/E)2. See also Figure S1.

Despite their origins as specialized forms of Pol II, the CTDs of the Pol IV and Pol V largest subunits share no sequence similarity with the CTD of the Pol II largest subunit, NRPB1 (Pontier et al. 2005; Haag and Pikaard 2011; Huang et al. 2015; Trujillo et al. 2016). Pol II largest subunit CTDs consist of repeats of the heptad consensus sequence, YSPTSPS, with varying numbers present in different species (e.g. 39 in Arabidopsis thaliana). These repeats can be phosphorylated in multiple patterns and mediate and numerous aspects of Pol II function, including polymerase recruitment, transcriptional activation, transcript elongation, termination, and RNA processing (reviewed in:Zaborowska et al. 2016).Whether the complex NRPE1 CTD mediates analogous Pol V functions is unclear. To functionally dissect the NRPE1 CTD, we engineered a series of NRPE1 transgenes with targeted CTD deletions and then tested their function. We show that the CTD is dispensable for Pol V transcription *in vitro* but is critical for Pol V transcript abundance and cytosine methylation *in vivo.* The biological activity of the CTD is explained primarily by the DeCL subdomain. However, the QS, 17aa repeat and linker subdomains are needed at loci that tend to be highly methylated and dependent on proteins Known to interact with Pol V or its transcripts. We show that RRP6L1, a paralog of the yeast ribonuclease Rrp6p, interacts with the QS and DeCL subdomains. Recombinant RRP6L1 produced in insect cells has 3'->5' exoribonuclease activity that is specific for single-stranded RNA, and is inhibited by RNA secondary structure. *In vivo*, our evidence indicates that RRP6L1 trims the 3' ends of Pol V transcripts sliced by AGO4, without completely degrading the RNAs, consistent with a prior study indicating, paradoxically, that RRP6L1 appears to stabilize Pol V transcripts associated with chromatin (Zhang et al., 2014). We propose a model whereby sliced transcripts engaged by RRP6L1 that is paused or slowed at regions of RNA secondary structure might increase the dwell time of AGO4-cleaved Pol V transcripts at their sites of synthesis, allowing these processed RNAs to contribute to RNA-directed DNA methylation.

## Results

### The NRPE1 CTD is required for Pol V-dependent RdDM

We engineered deletions affecting four CTD subdomains of the NRPE1 protein of *A. thaliana* ecotype Col-0: the linker subdomain (amino acid (aa) positions 1251-1426); the repeat subdomain (aa 1426-1651), containing ten complete, and two degenerate, repeats of a 17 aa consensus sequence, DKKNSETESGPAAWGSW; the DeCL subdomain (aa 1736-1851); and the QS (glutamine-serine)-rich subdomain at the very C-terminus (aa 1851-1976) (Figure 1A). Within the CTD are seventeen WG (Tryptophan-Glycine), one GWG, and one GW motif. Twelve of these Ago-hook motifs occur within the 17 aa repeat subdomain (aa 1426-1651) and mediate interactions with AGO4 (El-Shami et al. 2007).

Transgenes expressing the deleted NRPE1 proteins were tested for their ability to rescue an *nrpe1-11* null mutant, in like a full-length, un-mutated *NRPE1* transgene that is able to rescue the mutant. One set of recombinant NRPE1 proteins was sequentially deleted from the C-terminus (Δ1851-1976, Δ1736-1976, Δ1651-1976, Δ1566-1976, Δ1426-1976, and Δ1251-1976). Three additional transgenes were engineered to have internal CTD deletions: Δ1251-1426 (deleting the linker subdomain), Δ1426-1651 (deleting all 17aa repeats), and Δ1251-1651 (deleting the linker and repeat subdomains). We also tested a construct, NRPE1∆SD, engineered by the Lagrange lab (El-Shami et al. 2007), which has both the repeat and QS-rich subdomains deleted (Δ1411-1707 and Δ1875-1976) (Figure 1B). The transgenes were expressed from the native NRPE1 promoter and each recombinant protein (except NRPE1∆SD) was engineered to have a C-terminal HA epitope tag. Immunoprecipitation using anti-HA resin showed that the recombinant proteins are expressed at similar levels, displaying doublet bands typical of NRPE1 until linker sequences between 1251 and 1426 are deleted (Figure 1C-D). The Pol V second-largest subunit, NRP(D/E)2, co-immunoprecipitates with all of the recombinant NRPE1 proteins, indicating that polymerase assembly is not disrupted by the CTD deletions (Figure 1C-D). Moreover, full length NRPE1 and NRPE1 Δ1251-1976 (full CTD deletion) both localize to the nucleus and yield signals consistent with association with chromatin and Cajal bodies, similar to native NRPE1 (Figure S1).

To test CTD deletions for Pol V function, we first examined DNA methylation at known Pol V target loci using Chop-PCR (Figure 2A). In this method, genomic DNA is first digested (chopped) using a methylation sensitive restriction endonuclease prior to conducting PCR with primers flanking the endonuclease recognition site. If the site is methylated, it is not cut, and PCR amplification occurs. If the site is unmethylated, the DNA is cut and PCR fails. DNA was digested with *Hae*III or *Alu*I, whose cutting is blocked by CHH methylation (where H is a nucleotide other than G), a hallmark of Pol V-dependent RdDM. All loci yielded PCR amplification products using DNA of wild-type Col-0, indicating full methylation, but not using DNA of the *nrpe1-11* mutant, in which RdDM is lost (Figure 2A, S2A). The full-length *NRPE1* transgene, and NRPE1 missing only the QS-rich domain (Δ1851-1976) both restored methylation to wild-type levels (Figure 2A, S2A). However, deletion of the DeCL domain (constructs Δ1736-1976, Δ1651-1976) impaired restoration of DNA methylation at most loci (Figure 2A, S2A), and also prevented restoration of *AtSN1* transposon silencing (Figure S2B) or restoration of siRNA levels at *AtCopia* or *45S* rRNA gene loci (Figure S2C). However, for some loci, such as *IGN5A* and *IGN26*, deletion of additional CTD subdomains was needed to reduce Chop-PCR signals to *nrpe1-11* mutant levels and, remarkably, for methylation of the site assayed at the *soloLTR* locus, the CTD is dispensable (Figure 2A).

**Figure 2.**
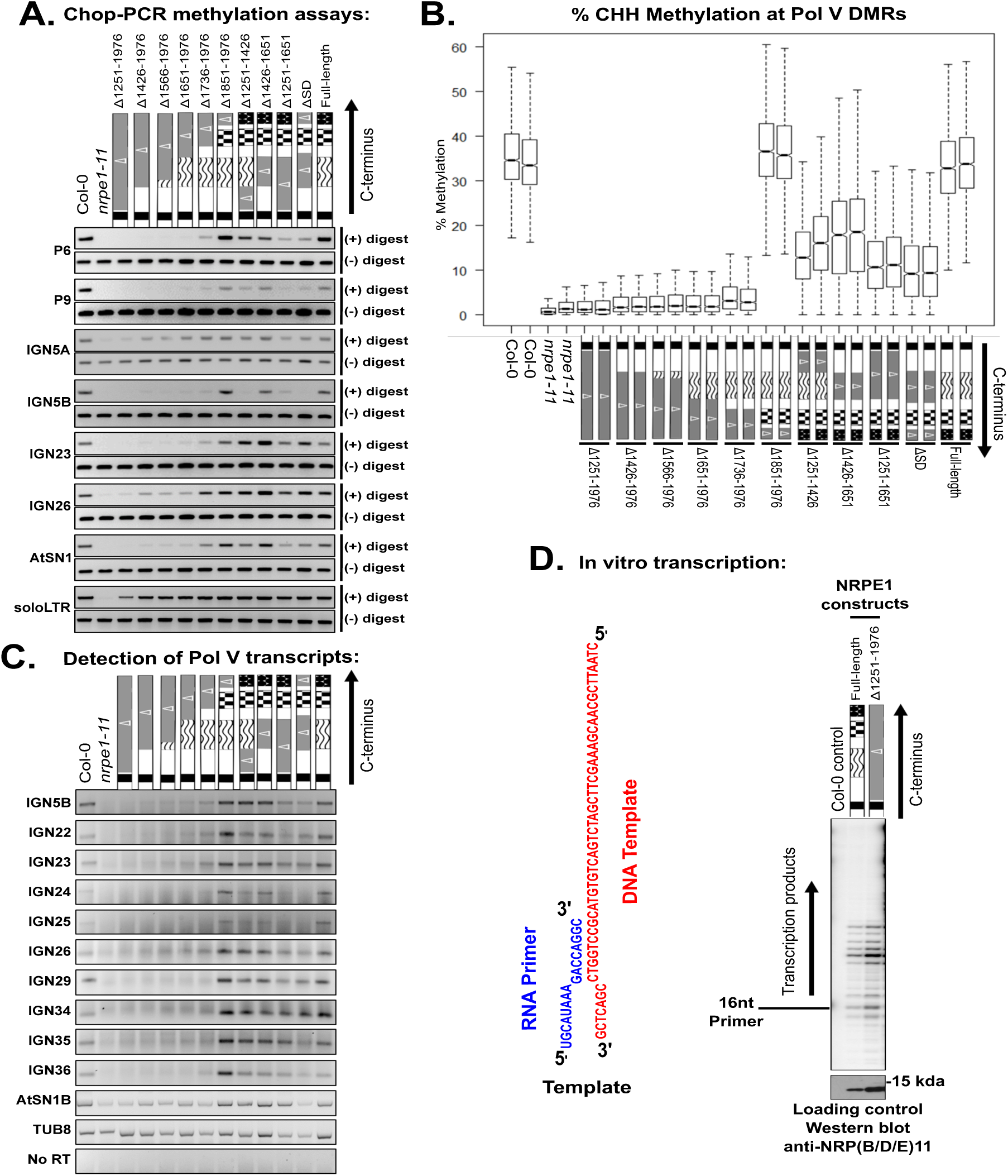
Effects of NRPE1 CTD deletions on CHH cytosine methylation and Pol V transcript abundance. Recombinant forms of NRPE1 were expressed in the *nrpe1-11* mutant background and tested for their ability to restore wild-type (ecotype Col-0) levels of methylation or Pol V transcript abundance. **A.** Chop-PCR assays testing methylation at *Alu*I or *Hae*III restriction endonuclease recognition sites at loci subjected to Pol V-dependent RNA-directed DNA methylation (RdDM). Transgenic lines expressing full-length NRPE1, or NRPE1 proteins bearing the indicated CTD deletions (shaded gray and denoted with the Greek symbol, Δ) are compared to wild-type (Col-0) and *nrpe1-11* controls. IGN5A and IGN5B are different PCR amplicon intervals in the same regions, with A being intergenic and B corresponding to sequences internal to a LINE element methylated and silenced by RdDM (seeWierzbicki et al, 2008). **B.** Box plots displaying overall cytosine methylation levels, determined by whole-genome bisulfite sequencing, within CHH motifs of Pol V-dependent differentially methylated regions (DMRs). Data for independent replicates conducted for each genotype are shown. **C.** RT-PCR detection of RNAs at ten Pol V-transcribed intergenic loci (IGN loci) and retrotransposon family, AtSN1 (PCR interval B; seeWierzbicki et al, 2008). *TUB8* (Tubulin), transcribed by Pol II, serves as a Pol V-independent control. **D.** Full length NRPE1 and NRPE1 missing the entire CTD have similar transcription activities, *in vitro.* NRPE1 proteins were immunoprecipitated from cell-free extracts of transgenic plants by virtue of C-terminal HA epitope tags. An immunoprecipitation fraction of non-transgenic Col-0 cell-free extract serves as a negative control. Relative protein input levels were compared by immunoblotting using anti-NRP(B/D/E)11 to detect the 11^th^ subunit, common to Pols II, IV and V. The NRPE1 fractions were incubated with a DNA oligonucleotide template hybridized to a RNA primer. Transcription products are labeled by virtue of incorporation of ^32^P-CTP and visualized by phosphorimaging (Haag et al, 2012). See also Figures S2-S3, and Tables S1-S2.

We next tested the effect of internal deletions eliminating subdomains N-terminal to the DeCL domain. NRPE1 proteins lacking the linker domain (Δ1251-1426) or the 17 aa repeat domain (Δ1426- 1651) partially restored DNA methylation, transcriptional silencing, and Pol V-dependent siRNA levels in the *nrpe1* mutant background, but did so in a locus-dependent manner (Figure 2A, Figure S2). NRPE1 proteins missing two CTD subdomains, namely Δ1251-1651 (linker and repeat domains deleted) and ΔSD (repeat and QS domains deleted), also did not fully rescue *nrpe1-11* methylation defects, with the degree of rescue being reduced compared to NRPE1 proteins deleted for either subdomain alone (Figure 2A, Figure S2).

### The DeCL subdomain accounts for most CTD activity

To assess how Pol V CTD subdomains impact DNA methylation genome-wide, we conducted whole genome bisulfite sequencing of DNA from wild-type, *nrpe1-11*, and plants expressing each CTD deletion construct in the *nrpe1* mutant background, with two biological replicates for each genotype (see Table S1 for sequencing statistics). In *nrpe1-11* plants, compared to wild-type, CHH methylation was lost from 2,259 differentially methylated regions (DMRs; see Methods) (Figure 2B, Table S2). Expressing full-length NRPE1, or NRPE1 missing only the QS domain (Δ1851-1976), restored CHH methylation at ~99% of these DMRs (Figure 2B), leaving only 16 or 30 DMRs, respectively, un-rescued (Figure S3A, S3G, Table S2). In contrast, in lines expressing NRPE1 missing the DeCL and QS domains (Δ1736-1976, Δ1651-1976, Δ1566-1976, Δ1426-1976, and Δ1251-1976), nearly 90% (~2,000) of the DMRs failed to regain methylation (Figure 2B, Figure S3 B-F, Table S2). NRPE1 proteins missing the linker or 17 aa repeat CTD subdomains restored methylation at 40-60 % of the DMRs (Figure 2B). Specifically, 922 DMRs that were demethylated in the *nrpe1* mutant remained demethylated in plants expressing NRPE1 missing the linker subdomain (Δ1251-1426), 685 demethylated DMRs persist when the 17 aa repeat subdomain (Δ1426-1651) is deleted, 1,246 demethylated DMRs persist when the linker and 17 aa repeat subdomains are deleted in tandem (Δ1251-1651 protein), and 1,421 demethylated DMRs persist for the ΔSD (repeat plus QS domain) protein (Figure S3H-I, Table S2).

### The NRPE1 CTD is required for Pol V transcript detection *in vivo* but not enzymatic activity *in vitro*

Pol V-dependent RNAs can be detected at RdDM target loci by RT-PCR (Wierzbicki et al. 2008; Wierzbicki et al. 2012), allowing us to test whether NRPE1 CTD deletion mutants that fail to rescue Pol V-dependent CHH methylation have reduced Pol V transcript levels *in vivo* (Figure 2C). Indeed, Pol V transcript levels are lowest in CTD deletion mutants in which CHH methylation is most reduced, with only trace (or background) levels of transcripts detected for NRPE1 proteins missing the DeCL domain. Deletion of the linker and 17 aa repeat domains affected Pol V transcript levels in a locus-dependent manner, whereas deletion of the QS domain had no effect at any of the loci tested (Figure 2C).

The severe loss of Pol V transcripts observed *in vivo* when the full CTD is deleted led us to test whether the Pol V CTD is required for the intrinsic RNA polymerase activity of Pol V. To do so, we conducted *in vitro* transcription using an assay in which an RNA primer annealed to a DNA template is elongated in a template-dependent manner (Haag et al. 2012). No difference was detected comparing full-length NRPE1 to NRPE1 missing the entire CTD (Δ1251-1976) (Figure 2D), indicating that the CTD is not required for Pol V's core catalytic activity. This suggests that CTD-effects on Pol V transcript abundance *in vivo* reflect functions required in a chromosomal or cellular context, such as target site recruitment, transcription initiation/elongation, or transcript stability.

### The full length NRPE1 CTD is required for RdDM at a subset of highly methylated loci

We examined the influence of CTD subdomains on CHH methylation of individual Pol V-dependent DMR loci using clustering analysis. Plants expressing full-length NRPE1, or NRPE1 missing just the QS domain (Δ1851-1976), display methylation profiles similar to wild-type plants (Figure 3A). In contrast, plants expressing NRPE1 lacking the DeCL domain (Δ1736-1976, Δ1651-1976, Δ1566-1976, Δ1426-1976, and Δ1251-1976) have methylation profiles similar to the *nrpe1-11* null mutant (Figure 3A). Profiles for NRPE1 proteins lacking the linker domain (Δ1251-1426), 17 aa repeat subdomain (Δ1426- 1651), linker plus 17 aa repeat subdomains (Δ1251-1651), or 17 aa repeat plus QS subdomain (∆SD), showed intermediate CHH methylation levels and affected overlapping subsets of Pol V-dependent loci (Figure 3A).

**Figure 3.**
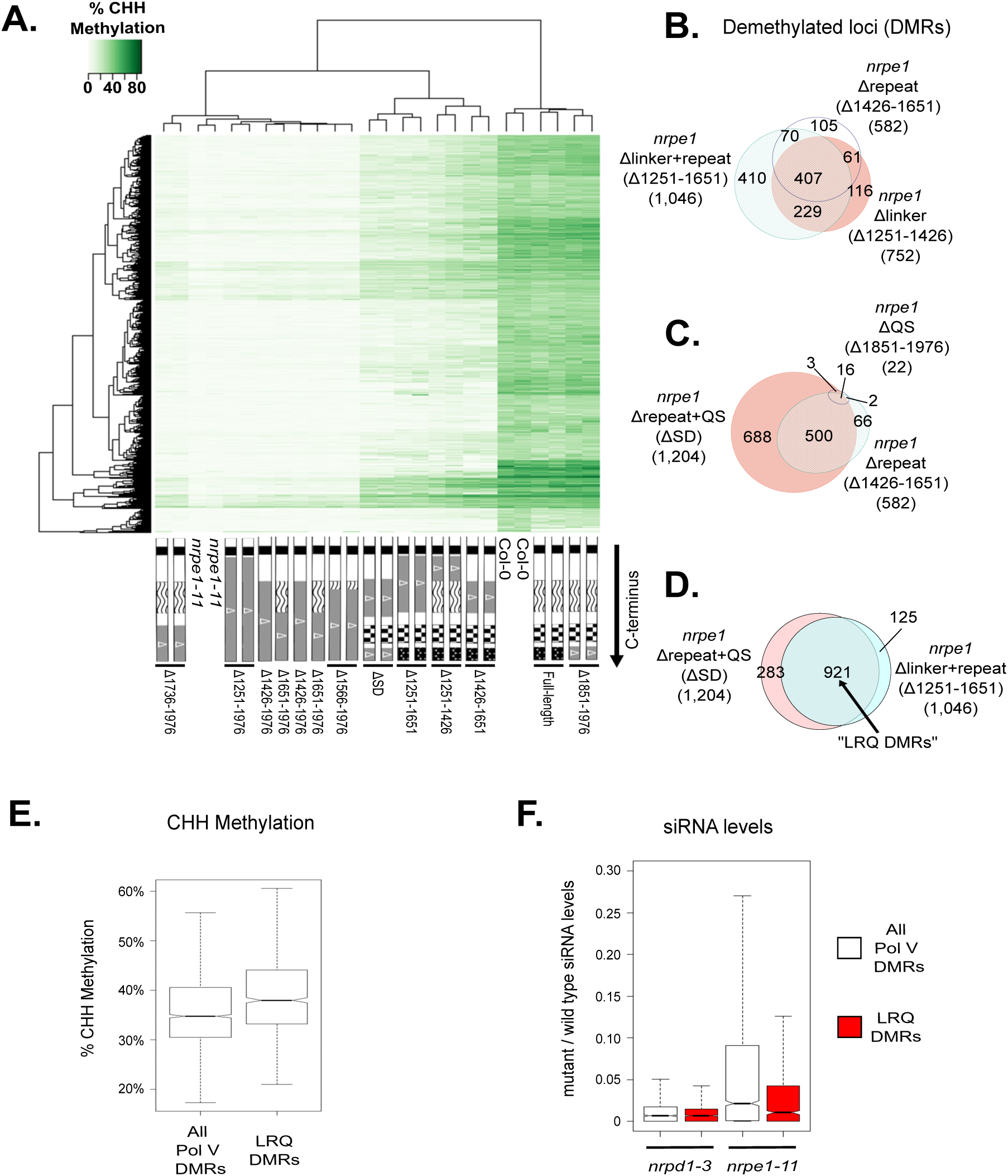
Functional relationships among Pol V CTD subdomains. **A.** Heatmap of % CHH methylation levels, depicted in shades of green, within Pol V DMRs for CTD deletion mutants, wild-type (Col-0) and *nrpe1-11* controls, arranged via clustering analysis. **B-D.** Venn diagrams showing subsets of the 2,259 total Pol V CHH DMRs that remain significantly hypo-methylated (un-rescued) when the indicated CTD mutants are expressed in the *nrpe1* mutant background. *Nrpe1* Δ1251-1651 and *nrpe1* ΔSD in (D) collectively represent deletions in the linker, repeat, and QS subdomains, thus the subset of DMRs un-rescued by both constructs are referred to as LRQ DMRs. **E.** Box plot of % CHH methylation in wild type plants within all Pol V CHH DMRs, compared to the 921 LRQ DMRs defined in panel D. **F.** Box plot showing the ratio of siRNA levels (in reads per million; RPM) in *nrpd1-3* and *nrpe1-11* mutants relative to wild-type Col-0. Total Pol V DMRs (white) are compared to the 921 LRQ DMRs defined in panel D (red). See also Table S2.

Comparing Pol V-dependent DMRs that fail to regain methylation when NRPE1 is missing either the linker subdomain (Δ1251-1426; 752 DMRs) or 17 aa repeat subdomain (Δ1426-1651; 582 DMRs) revealed an overlap of 468 DMRs (Figure 3B). If the linker and 17 aa repeat subdomains are both deleted (Δ1251-1651), the effect is more severe, such that nearly half of all Pol V-dependent DMRs (1,046 out of 2,259) remain significantly hypo-methylated, including 410 DMRs that were not affected by deletion of either subdomain alone (Figure 3B), suggesting partially redundant functions for these adjacent subdomains of the CTD.

We next compared the overlap between Pol V-dependent DMRs that remain significantly hypo-methylated when NRPE1 lacks the QS domain (Δ1851-1976) or the repeat domain (Δ1426-1651) (Figure 3C). Only 1% (22) of the 2,259 total Pol V DMRs remain hypo-methylated when NRPE1 lacking the QS domain is expressed in the *nrpe1* mutant. Eighteen of these DMRs are among the 582 DMRs that remain hypo-methylated when the 17 aa repeat domain is deleted (Δ1426-1651) (Figure 3C). Interestingly, deleting both the QS and 17 aa repeat subdomains (construct ΔSD) is substantially more deleterious that deleting either domain alone, such that 1,204 DMRs are not rescued, suggesting a functional interaction between these domains (Figure 3C).

We compared the effect of deleting the linker plus 17 aa repeat subdomains (Δ1251-1651) to ΔSD, in which the 17aa repeat and QS subdomains are deleted. 921 hypo-methylated regions overlap among the 1,204 (∆SD) and 1,046 (∆1251-1651)-affected DMRs, accounting for approximately half of the 2,259 Pol V-dependent DMRs (Figure 3D). Because the Δ1251-1651 and ΔSD forms of NRPE1 both have the critical DeCL subdomain, but are deleted for the linker (L), repeat (R), and QS (Q) subdomains, we refer to the 921 DMRs affected by both constructs as LRQ DMRs for the remainder of the manuscript (Table S2). LRQ DMRs have at least two notable characteristics. One is that their methylation level in wild-type Col-0 is higher, on average, than total Pol V-dependent DMRs (Figure 3E). Second, they differ with respect to siRNA levels. Pol IV is required for the biogenesis of virtually all 24nt siRNAs (Pontier et al. 2005; Zhang et al. 2007; Mosher et al. 2008), such that dividing the number of 24nt siRNAs detected in Pol IV (*nrpd1-3*) mutants by the number in wild type (Col-0) yields an average value near zero at both LRQ DMRs and total Pol V DMRs (Figure 3F, Table S3). In contrast, Pol V affects siRNA production at only a subset of loci (Pontier et al. 2005; Mosher et al. 2008). At LRQ DMRs, siRNA levels in *nrpe1-11* mutants are substantially lower than for Pol V DMRs as a whole, indicating that siRNA levels at LRQ DMRs have a greater dependency on Pol V (Figure 3F, Table S3).

### LRQ DMRs correlate with loci whose methylation is similarly dependent on Pol V transcript binding proteins

Comparing CHH methylation profiles of CTD mutants and mutants acting in Pol V-dependent steps of the RdDM pathway revealed intriguing clustering relationships (Figure 4A). Mutants with severe methylation defects, clustering with *nrpe1-11*, include NRPE1 with the full CTD deleted (Δ1251-1976), NRPE1 missing the DeCL subdomain (Δ1736-1976) and several mutants defective for proteins implicated in Pol V recruitment to target sites, including *drd1, dms3 and suvh2 suvh9* (Wierzbicki et al. 2008; Wierzbicki et al. 2009; Zhong et al. 2012; Johnson et al. 2014; Liu et al. 2014; Jing et al. 2016) (Figure 4A, Table S2). Mutants with less severe effects on methylation, resembling CTD deletion mutants lacking the linker, 17 aa repeat, or QS subdomains, include mutants defective for proteins that interact with Pol V transcripts and/or Pol V transcription elongation complexes, including IDN2-IDP complex (*idn2 idnl1 idnl2*)*, spt5L (*also known as *ktf1)*, and *rrp6L1*, (Ausin et al. 2009; Bies-Etheve et al. 2009; He et al. 2009; Rowley et al. 2011; Zhang et al. 2012; Kollen et al. 2015) (Figure 4A, Table S2). The IDN2-IDP complex binds Pol V transcripts prior to recruitment of the *de novo* DNA methyltransferase, DRM2 (Ausin et al. 2009; Ausin et al. 2012; Bohmdorfer et al. 2014). SPT5L is a paralog of the Pol II transcription elongation factor, SPT5, and can co-immunoprecipitate (at low levels) with Pol V, bind Pol V transcripts and physically interact with AGO4 and AGO6 via extensive ago-hook motifs (Bies-Etheve et al. 2009; He et al. 2009; Huang et al. 2009; Rowley et al. 2011). RRP6L1 binds Pol V transcripts and has been implicated in maintaining these RNAs in chromatin (Zhang et al. 2014). The grouping of these different mutants by clustering analyses was independently validated by comparing statistically significant hypo-DMRs in each mutant line to LRQ DMRs, confirming a high degree of overlap (Figure 4B-D, Table S2).

**Figure 4.**
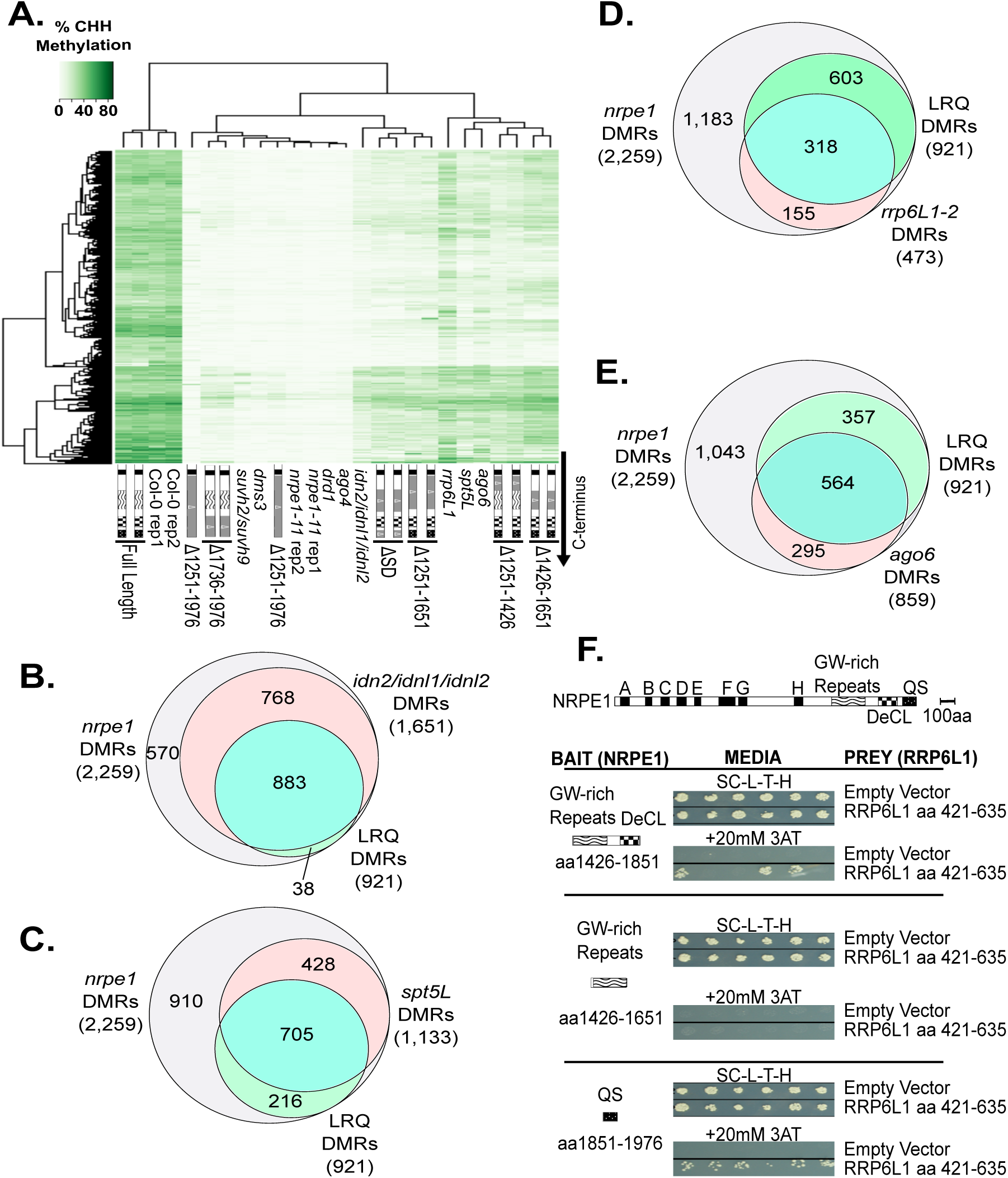
Methylation defects resulting from Pol V CTD deletions resemble defects observed in other RdDM mutants. **A.** Heatmap and clustering analysis of % CHH methylation at Pol V DMRs, comparing CTD mutants, other RdDM pathway mutants (*suvh2 suvh9, dms3, drd1, ago4, ago6, idn2 idnl1 idnl2, rrp6L1, spt5*), and wild-type (Col-0) and *nrpe1-11* controls. **B-E.** Venn diagrams comparing total Pol V CHH DMRs, and LRQ DMRs in combination with DMRs dependent on *idn2/idnl1/idnl2* (**B**), *spt5L* (**C**), *rrp6L1* (**D**), or *ago6* (**E**). **F.** Yeast two hybrid (Y2H) interaction tests for Pol V CTD subdomains (“Bait”) and RRP6L1 aa 426-635 (“Prey”). Each bait was also tested against empty vector (pDEST22) controls. SC-L-T-H refers to media lacking leucine, tryptophan and histidine (top panel). Bait-prey interaction allows growth in the presence of 20 mM E-Amino-1, 2, 4 Triazol (3-AT), a HIS3 inhibitor (bottom panel). See also Figure S4 and Tables S1 and S2.

AGO4 and AGO6 interact with Pol V transcripts as well as AGO-hook motifs of the CTD 17 aa repeat subdomain (aa 1426-1651) (El-Shami et al. 2007; Wierzbicki et al. 2009; Rowley et al. 2011; Pontier et al. 2012). Thus, one might predict that methylation defects resulting from targeted deletion of the 17 aa repeat subdomain might be phenocopied by *ago4* or *ago6* mutants. Interestingly, DMRs affected by *ago6* and the 17 aa repeat subdomain fit this expectation and show substantial overlap (Figure 4A, E). However, *ago4* mutants exhibit a much more severe loss of CHH methylation (Stroud et al. 2013) than NRPE1 lacking the 17 aa repeat subdomain, clustering with the *nrpe1-11*, full CTD, or DeCL subdomain deletions (Figure 4A). This finding is consistent with evidence that AGO4 recruitment entails more than 17 aa repeat subdomain interactions, including interactions with Pol V transcripts and SPT5L (El-Shami et al. 2007; Wierzbicki et al. 2009; Pontier et al. 2012; Lahmy et al. 2016; Wendte and Pikaard 2016).

### The exonuclease, RRP6L1, interacts with the CTD and trims Pol V transcripts

We conducted a yeast two hybrid (Y2H) screen for *A. thaliana* proteins that interact with the Pol V CTD. The full CTD could not be used as bait due to auto-activation of the reporter gene, leading us to test individual domains. Only one confirmed interactor was identified, using the QS subdomain (aa 1851-1976) as bait. This interactor is RRP6L1 (see Figure 4A), previously identified by the Zhu lab (Zhang et al., 2014) as a protein affecting RdDM at a subset of loci and stabilizing Pol V transcripts in the context of chromatin, an apparent paradox given that the yeast paralog, Rrp6p is a 3’ to 5’ exoribonuclease (Lange et al. 2008; Fox and Mosley 2016) (Figure 4F). Confirming the Y2H results, full-length RRP6L1 expressed in insect cells physically interacts with a recombinant QS subdomain polypeptide *in vitro* (Figure S4A-B). Additional Y2H tests showed that RRP6L1 also interacts with a 17 aa repeat plus DeCL subdomain polypeptide (NRPE1 amino acids 1426-1851), but not the 17 aa repeat subdomain alone (aa 1426-1651). Collectively, these results suggest that RRP6L1 can interact with both the DeCL and QS subdomains of the Pol V CTD (Figure 4F).

One might expect that if RRP6L1 degrades Pol V transcripts, Pol V transcript levels would increase in *rrp6L1* mutants, but this is not the case (Figure 5A). This led us to test the possibility that RRP6L1 is non-functional as a nuclease. However, RRP6L1 expressed in insect cells (Figure S4A) exhibits exonuclease activity, acting on single-stranded RNA that has a 3' hydroxyl group, but not singlestranded RNA with a 3’ phosphate, single-stranded DNA, single-stranded RNA with secondary structure, or double-stranded RNA with 100% basepairing (Figure 5B).

**Figure 5.**
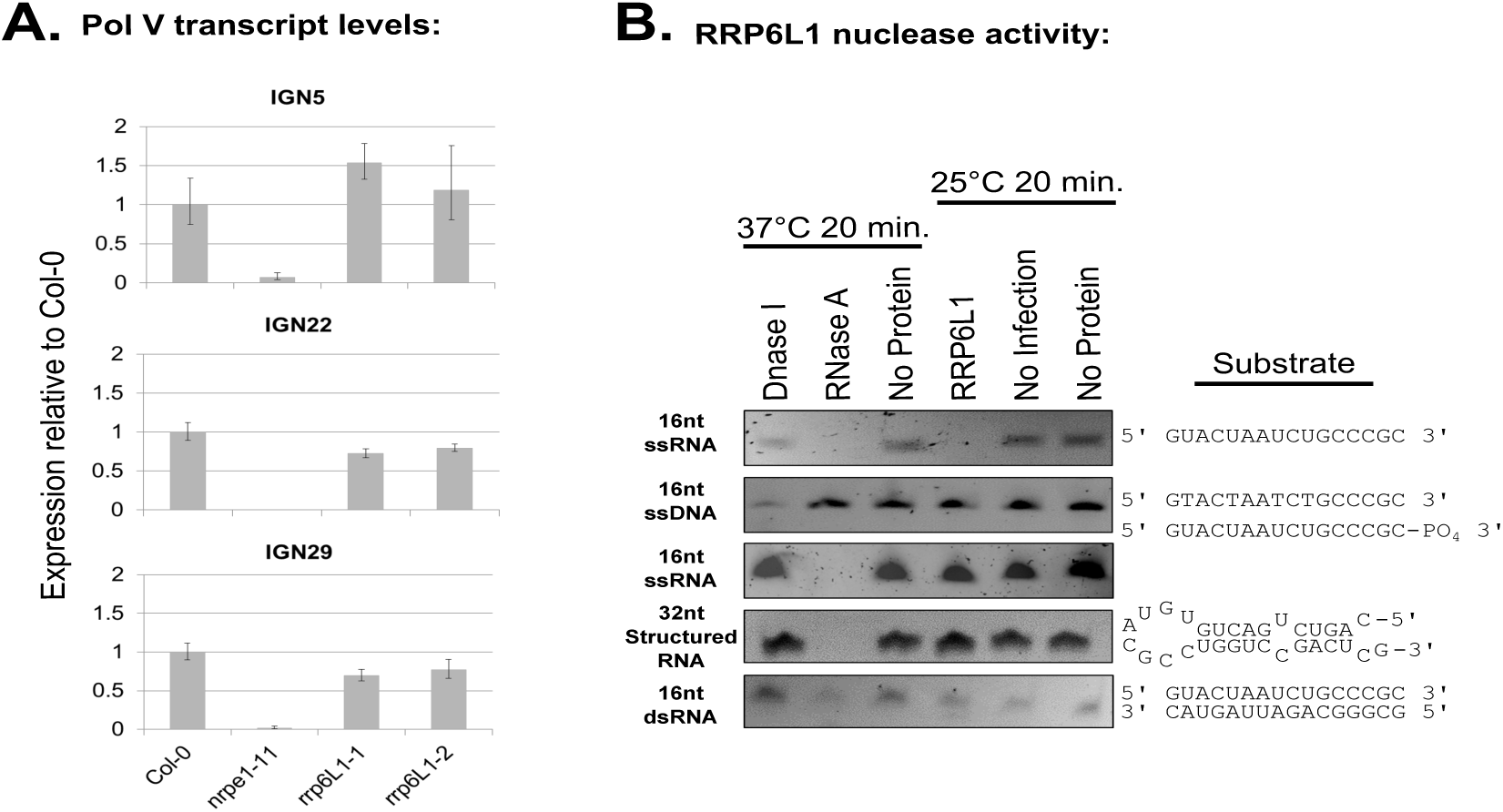
The CTD-interactor, RRP6L1 is a 3’ to 5’ exonuclease **A.** Quantitative RT-PCR of Pol V-transcript levels in the wild-type (Col-0), *nrpe1-11* or *rrp6L1* mutants at three IGN loci. Histograms show ratios of ΔCt: (Ct - Ct_*Actin2*_) values in the indicated genotypes relative to Col-0. Error bars represent the propagated standard error of the mean for 3 technical replicates. **B.** SYBR gold-stained polyacrylamide gels of RNA or DNA substrates following incubations with DNase I, RNase A, RRP6L1 expressed in insect cells from a baculovirus vector, or buffer (no protein). No infection controls are cell-free extracts of insect cells not infected with baculovirus but otherwise treated the same as for RRP6L1 extracts. See also Figure S4.

Our experimental evidence shows that RRP6L1 is a functional exonuclease that interacts with the Pol V CTD, yet the Zhu lab has shown that RRP6L1 interacts with Pol V transcripts and somehow stabilizes the RNAs in the context of chromatin (Zhang et al., 2014). A potential solution to this paradox might be that RRP6L1 does not completely degrade Pol V transcripts, but merely trims their 3’ ends, analogous to yeast Rrp6p's trimming of several nuclear RNA species (Fox and Mosley 2016). To test this hypothesis, we devised a modified 3’ RACE procedure to amplify and sequence the 3' ends of RNAs at known Pol V-transcribed loci (Table S4). At the *IGN5B* and *IGN25* loci, where CHH methylation is RRP6L1-dependent, the median length of transcript 3' ends increased in *rrp6L1-1* and *rrp6L1-2* mutants, relative to wild-type (Figures 6A and 6B). In contrast, at *IGN17, IGN23, IGN29* and *IGN35*, where RRP6L1 has no effect on CHH methylation, transcript 3' end lengths were unaffected, or slightly shorter, in *rrp6L1* mutants (Figure 6C-D, Figure S5). Overall, these results suggest that at loci where CHH methylation is dependent on RRP6L1, the exonuclease trims the 3’ ends of Pol V transcripts.

**Figure 6.**
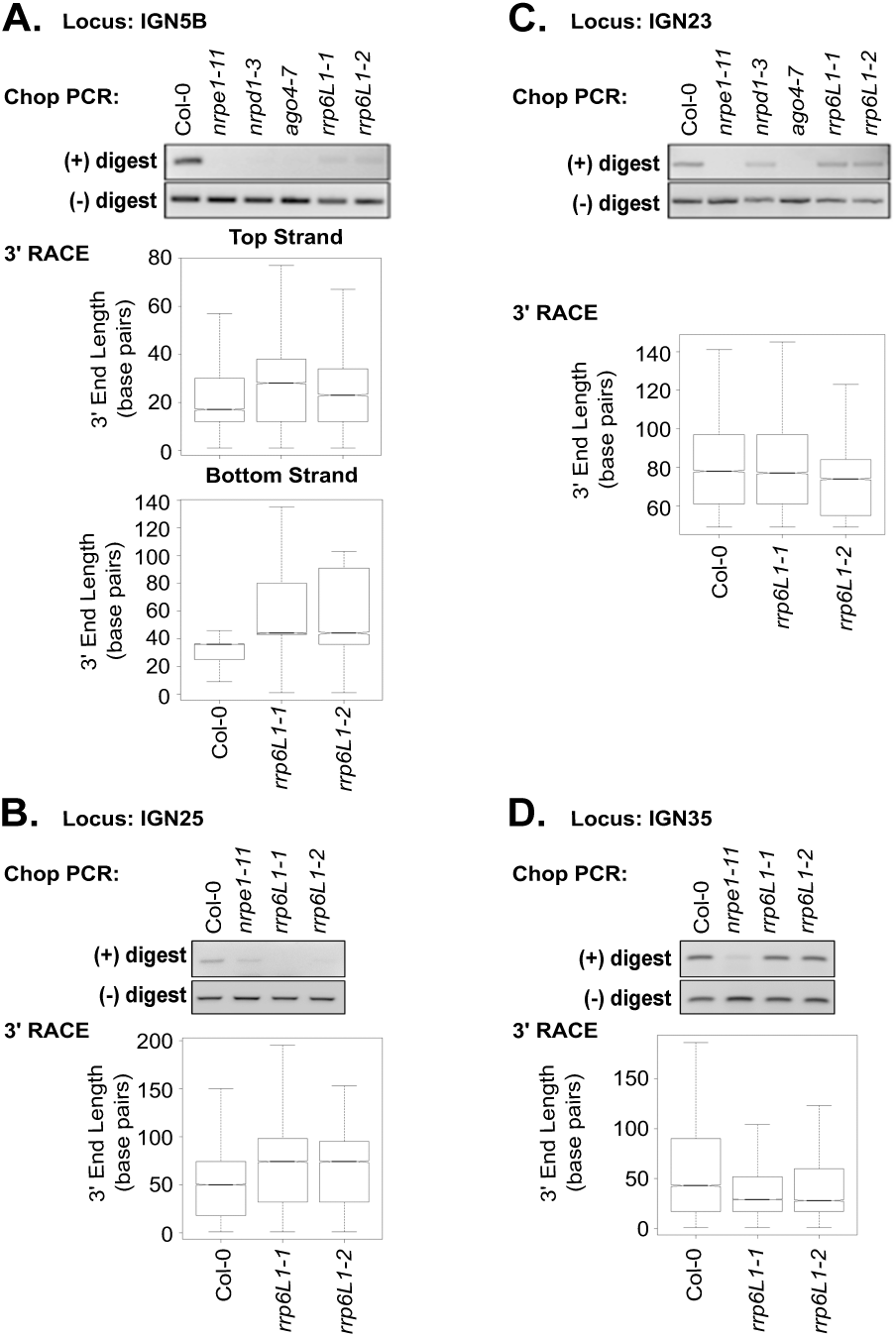
Evidence for RRP6L1 trimming of Pol V transcript 3’ ends. Chop-PCR and 3’ RACE results for Pol V transcribed loci: **A.** *IGN5*, **B.** *IGN25*, **C.** *IGN23*, **D.** *IGN35.* Chop PCR was conducted with the enzymes *Hae*III or *Alu*I, both of which are inhibited by CHH methylation. Box plots show 3’ end lengths in basepairs, measured from the internal gene specific primer used in the 3’ RACE reactions to the ends of the RNAs. See also Figure S5 and Table S4.

As an independent test for longer than normal RNAs at RRP6L1-dependent loci, we examined the IGN5 locus, where Pol V-dependent transcripts can be readily detected by RT-PCR using a primer pair defining what is termed amplicon 1 in Figure S5C. Interestingly, amplicon 1 RT-PCR signals are elevated in mutants for Pol IV (*nrpd1*) or *ago4*, compared to wild-type, suggesting that Pol V transcript levels increase in the absence of AGO4 cleavage directed by Pol IV-dependent siRNAs. The RT-PCR signals similarly increase in the *rrp6L1-1* mutant, and are not further increased in an *ago4 rrp6L1* double mutant, suggesting functions in the same pathway. However, the signals are lost in *rrp6L1 nrpe1* double mutants, confirming that Pol V transcripts are being monitored (Figure S5C).

Moving one PCR primer an additional 58 bp in the 3’ direction defines a larger amplicon, amplicon 2, for which corresponding RNAs are not detected in wild-type. However, amplicon 2 transcripts are detected above background levels in *nrpd1, ago4*, or *rrp6L1* mutants, do not increase in *ago4 rrp6L1* double mutants, and are lost in *rrp6L1 nrpe1* double mutants (Figure S5C). Collectively, we interpret these RT-PCT results as evidence that RRP6L1 trims Pol V transcript 3' ends generated by siRNA-directed AGO4 slicing.

## Discussion

The CTD of NRPE1 is essential for Pol V function *in vivo*, but not for Pol V subunit assembly, nuclear localization or RNA polymerase activity *in vitro.* Thus, the reduced or undetectable levels of Pol V transcripts in full CTD or DeCL subdomain deletion mutants suggests an important role for the CTD in transcript production in context of chromatin, possibly affecting Pol V recruitment to target sites, initiation, elongation, or transcript stability. Consistent with this interpretation, Pol V transcripts and RdDM is similarly reduced in mutants for DRD1, a putative ATP-dependent DNA translocase, or DMS3, a protein related to the hinge domains of cohesins and condensins (Wierzbicki et al. 2008; Wierzbicki et al. 2009), that are components of a multi-protein complex (abbreviated DDR) needed for Pol V to be detected at target loci by chromatin immunoprecipitation (Law et al. 2010; Zhong et al. 2012). Likewise, the methylcytosine binding proteins SUVH2 and SUVH9 are also critical for Pol V recruitment to methylated target loci and interact with the DDR complex (Johnson et al. 2008; Liu et al. 2014). Importantly, the methylation profiles for *suvh2 suvh9* double mutants, *drd1, dms3*, and *nrpe1* mutants lacking the full CTD or DeCL subdomain are similar (see Figure 4A).

Deleting the DeCL subdomain has nearly the same effect as deleting the entire CTD. Other CTD subdomains are not as critical as the DeCL, but they significantly affect methylation at subsets of loci. Interestingly, methylated loci dependent on the linker, 17aa repeat or QS subdomains are similarly dependent on proteins that interact with Pol V or its transcripts, including SPT5L, the IDP complex (IDN2, IDNL1, IDNL2), AGO6, or RRP6L1, implicating these CTD subdomains in co-transcriptional steps of the RdDM process.

Repeated peptide sequences are present in all plant NRPE1 CTDs, yet their number and sequence varies widely, suggesting rapid evolution enabled by low selective pressure (Trujillo et al. 2016). In our NRPE1 deletion mutant that removed all 12 of the 17 aa repeats, methylation was still rescued at ~60% of Pol V-dependent loci, suggesting that these repeats are non-essential at most loci. The 17 aa repeats account for the majority of WG/GW Ago-hook motifs in the CTD, and are the major site of AGO4 interaction with Pol V (El-Shami et al. 2007). Interestingly, a study published while this manuscript was in preparation showed that restoring 17 aa repeats into the NRPE1∆SD construct rescues DNA methylation at affected loci, even upon changing the WG motifs to AG motifs, suggesting an unknown function for the repeats other than AGO binding (Lahmy et al. 2016).

The QS subdomain, located at the extreme C-terminus of NRPE1, is the least conserved feature of plant NRPE1 proteins, being present in *A. thaliana* but absent in other genera of the Brassicaceae family, including the closely related species, *A. lyrata.* Although the QS subdomain can be deleted without apparent consequence for Pol V function, its deletion has a synergistic effect when in combination with deletion of the 17 aa repeat subdomain. The QS and DeCL subdomains each interact with RRP6L1, suggesting that the QS subdomain may serve a recently evolved function that is at least partially redundant with the functions of other CTD subdomains.

Arabidopsis RRP6L1 was identified previously in a genetic screen for mutants disrupted in RdDM and found to stabilize Pol V transcript associations with chromatin, despite being a predicted exonuclease (Zhang et al., 2014). Our results provide several new insights, summarized in the model of Figure 7. First, our biochemical evidence indicates that RRP6L1 is an exonuclease that requires RNA with a free 3’ hydroxyl group, which would be present at the end of Pol V transcripts following termination and release, or would be present upon cleavage internally by an endonuclease, such as AGO4. The fact that Pol V transcript levels are similarly elevated in *ago4* and *rrp6L1* mutants, or mutants defective for siRNA biogenesis, is consistent with the latter hypothesis. Furthermore, Pol V transcripts exist in cells as a mixed population of species possessing either a 5’ triphosphate or 5’ monophosphate, indicative of both primary and sliced products (Wierzbicki et al. 2008; Wendte and Pikaard 2016). It is noteworthy that AGO4 and RRP6L1-interacting subdomains of the Pol V CTD are adjacent to one another, such that siRNA-guided AGO4 slicing (Qi et al. 2006) may be coupled to RRP6L1 engagement of resulting RNA 3' ends. *In vitro*, RRP6L1 exonuclease activity is inhibited by RNA secondary structure, suggesting that pausing at structured RNA regions might explain Zhang et al.'s finding that RRP6L1 helps retain Pol V transcripts in chromatin, which could promote engagement of the Pol V transcript-binding IDP complex and subsequent recruitment of the DNA methyltransferase, DRM2 (Figure 7).

**Figure 7.**
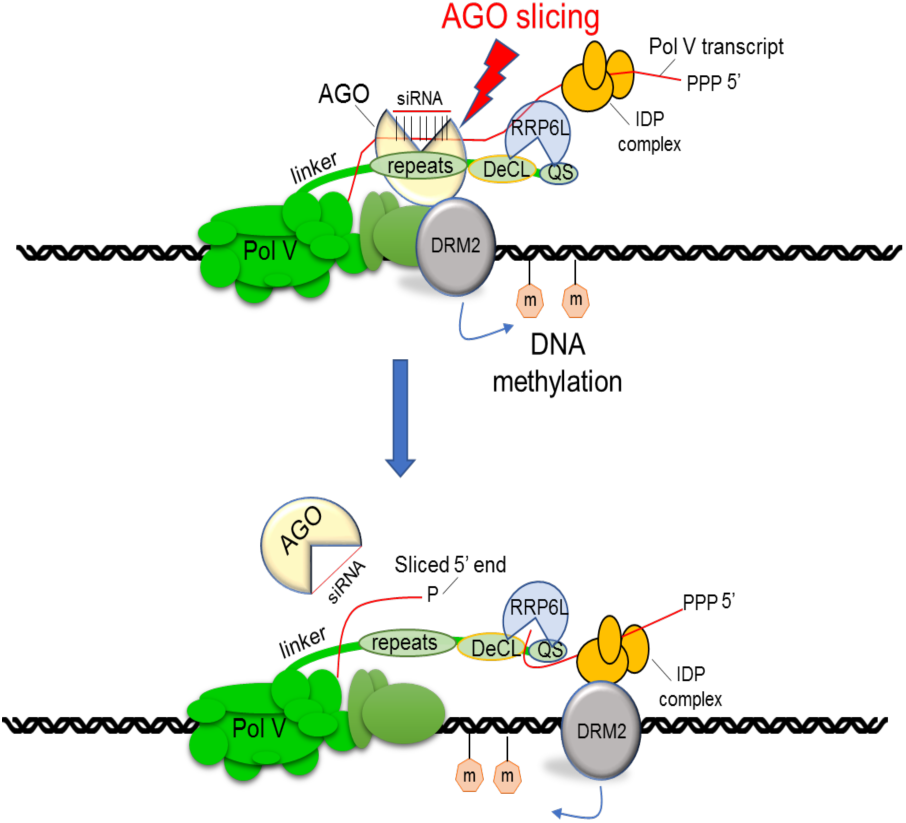
Model for CTD-mediated coordination of AGO4 transcript cleavage and RRP6L1 engagement of cleaved RNA 3' ends. AGO4 and RRP6L1 bind adjacent subdomains of the NRPE1 CTD, such that cotranscriptional slicing of Pol V transcripts by AGO4, guided by basepaired 24 nt siRNAs, may be coupled to RRP6L1 engagement of cleaved RNA 3’ ends. RRP6L1's trimming of Pol V transcripts, with pausing at sites of secondary structure, may facilitate RNA retention, allowing Pol V transcript-binding proteins, such as the IDP complex, to recruit the *de novo* cytosine methyltransferase, DRM2 to cleaved RNAs while Pol V transcription of nascent RNA continues.

The Pol V largest catalytic subunit gene, *NRPE1* clearly evolved from a duplicated copy of the ancestral Pol II subunit gene, *NRPB1*, but the 3' exons of the two genes are unrelated, such that the CTDs of the encoded proteins share no detectable sequence similarity. Pol II largest subunits in all eukaryotes have CTDs composed entirely of repeating units of a heptapeptide whose consensus is YSPTSPS. These heptad repeats can be phoshorylated in multiple patterns and play roles in virtually all aspects of Pol II activity, including recruitment to gene promoters, pre-initiation complex assembly, transcription initiation, transcript elongation, addition of 7-methyguanosine caps to mRNA 5' ends, mRNA processing, transcription termination and transcription-coupled chromatin modification (reviewed in:Zaborowska et al. 2016). Pol V and Pol IV lack the heptad repeats of Pol II, and the CTD of Pol V is complex, spanning ~700 amino acids and including simple repeats (QS), longer repeats (17 aa), and a subdomain related to a chloroplast protein family (DeCL). Instead of serving as interaction sites for proteins important for mRNA synthesis and processing, Pol V CTD subdomains mediate processes specific to RNA-directed silencing, including Argonaute and RRP6L interactions. Further understanding of the molecular details of CTD-mediated processes will likely help illuminate the full spectrum of Pol V functions.

## Materials and Methods

### Data availability

The deep sequencing data generated for this study have been deposited in NCBI's Gene Expression Omnibus (Edgar et al. 2002) and are accessible through GEO Series accession number GSE93360 (https://www.ncbi.nlm.nih.gov/geo/query/acc.cgi?acc=GSE93360).

Pre-publication reviewer access link:
https://www.ncbi.nlm.nih.qov/qeo/querv/acc.cgi?token=mhchwsocfnmdnqh&acc=GSE93360

### Plant material

*Arabidopsis thaliana* mutant lines, *nrpe1-11* (Salk 029919), *nrpd1-3* (Salk 128428), *rrp6L1-1* (Salk 004432), *rrp6L1-2* (Gabi 344G09) are available through the *Arabidopsis* resource center (TAIR) and have been described previously (Onodera et al. 2005; Pontier et al. 2005; Zhang et al. 2014). The *ago4-1* mutant is described in (Wierzbicki et al. 2009). Plants were grown in soil in long day conditions (16 hours light, 8 hours dark).

### Targeted CTD deletion construct generation and plant transformation

The pENTR-NRPE1 full-length genomic sequence with its endogenous promoter (Pontes et al. 2006) was recombined into pEarleyGate301 (Earley et al. 2006) to add a C-terminal HA tag. C-terminal domain deletions were obtained by using the pENTR-NRPE 1 genomic clone as a DNA template with reverse primers that truncated the 3’ end (see Table S5 for primer sequences). Pfu Ultra (Stratagene) was used to amplify the sequences. The PCR products were gel purified and cloned into pENTR-TOPO S/D (Invitrogen) before being recombined into pEarleyGate 301. Internal C-terminal domain deletions, *nrpe1* Δ1251-1426 and *nrpe1* Δ1251-1651, were obtained by the SLIM method (Chiu et al. 2004), and *nrpe1* Δ1426-1651 was obtained using Stratagene cloning (now StrataClone from Agilent), using appropriate primers (Table S5). pEarleyGate plasmids (10-50 ng) were transformed into *Agrobacterium tumefaciens* which was used to transform *nrpe1-11* plants using the floral dip method (Bechtold and Pelletier 1998; Clough and Bent 1998).

### Immunoprecipitation and western blot analysis

Protein was extracted from frozen leaf tissue (4.0 g) as described in (Pontes et al. 2006). Protein samples were subjected to SDS-PAGE run on Tris-glycine gels and transferred to nitrocellulose or PVDF membranes. Antibodies were diluted in TBST + 5% (w/v) nonfat dried milk as follows: 1:500 NRPD/E2, 1:500 anti-NRPB/D/E11, and 1:3,000 anti-HA-HRP. Anti-rabbit-HRP (Amersham or Santa Cruz Biotechnology) diluted 1:5,000 was used as secondary antibody for native antibodies. Rabbit antibodies to NRPB11/NRPD11/NRPE11 (AT3G52090) were generated by Sigma and purified using immobilized NRPB/D/E11 protein.

### Chop PCR

Chop PCR methylation analyses described in Figure 2A and Figure 6 were conducted using DNA extracted from 2.5 week old above ground plant tissues using the CTAB method (Murray and Thompson 1980). ~350 ng of DNA was double digested using the enzymes *Alu*I and *Hae*III (NEB) at 37°C for 3 hours. PCR of regions of interest was conducted using GoTAQ Green (Promega) and primers described in Table S5.

### RT-PCR

RNA for RT-PCR was extracted from ~2.5 week old above ground plant tissues. Semi-quantitative RT-PCR analyses shown in Figure 2C and S5C were conducted as described in (Wierzbicki et al. 2008).Quantitative real time PCR shown in Figure 5A was conducted as described in (Rowley et al. 2013). All primer sequences are in Table S5.

### In vitro transcription assays

In vitro transcription assays were conducted as described in (Haag et al. 2012) with the following changes. Polymerases were immunoprecipitated from 4 g frozen leaf tissue as described previously, except that 150 mM NaSO_4_ and 5mM MgSO_4_ were used in extraction buffer in place of NaCl and MgCl_2_, 25 μl HA resin was used for the IP, and NP-40 was not added to the wash buffer. Resin-associated polymerases were washed once with CB100 buffer (100 mM potassium acetate, 25 mM HEPES, pH 7.9, 20% glycerol, 0.1 mM EDTA, 0.5 mM DTT, 1 mM PMSF), resuspended in 50 μL CB100 buffer and supplemented with 50 μL 2x transcription reaction buffer (120 mM ammonium sulfate, 40 mM HEPES, pH 7.6, 20 mM magnesium sulfate, 20 μM zinc sulfate, 20% glycerol, 0.16 U/μL RNaseOUT, 20 mM DTT, 2 mM ATP, 2 mM UTP, 2 mM GTP, 0.08 mM CTP, 0.2 mCi/mL alpha ^32^P-CTP and 4 pmols of template. Transcription reactions were incubated for 60 minutes at room temperature on a rotating mixer, and stopped by addition of 50 mM EDTA and heating at 75 °C for five minutes. Reaction products were enriched using PERFORMA spin columns (EdgeBio) as per manufacturer’s protocol and precipitated using 1/10 volume of 3M sodium acetate, pH 5.2, 20 μg glycogen and 2 volumes isopropanol at −20°C overnight. Precipitated RNA was resuspended in 5 μL 2X RNA loading buffer (NEB), incubated at 70°C for 5 minutes and loaded on to 15% denaturing polyacrylamide gels. Gels were transferred to Whatman 3MM filter paper, dried under vacuum for 2 hours at 80 °C, and subjected to phosphorimaging.

Templates for *in vitro* transcription were generated by using equimolar amounts (10 mM each) of DNA template and the RNA primer were mixed in the annealing buffer containing 20 mM HEPES-KOH (pH 7.6) and 100 mM potassium acetate. The mixture was boiled in a water bath and slowly cooled to room temperature. Template sequences are in Table S5.

### Bisulfite sequencing

100 μg of DNA extracted from ~2.5 week old above ground plant tissues was prepared for Illumina sequencing using the TruSeq DNA methylation library prep kit according to the manufacturer’s instructions. Libraries were generated using an Illumina NextSeq instrument. Detailed procedures for mapping and analysis of bisulfite sequencing data are in the Supplemental Methods. Accession numbers for previously published bisulfite sequencing data analyzed are in Table S1.

### Small RNA sequencing

Small RNA sequencing for *nrpe1-11* was completed as described in (Blevins et al. 2015) by Fasteris SA (http://www.fasteris.com/). Detailed procedures for mapping and analysis of sRNA sequencing data are in the Supplemental Methods. Accession numbers for previously published sRNA data for Col-0 and *nrpd1-3* are SRR2075819 and SRR2505369, respectively.

### Yeast two hybrid analyses

Yeast two hybrid (Y2H) was performed by the Indiana University Yeast Two Hybrid Facility. Briefly, an uncut custom cDNA Library for *A. thaliana* in the entry vector, pENTR222 (Invitrogen) was cloned into the prey vector, pDEST22 using LR Clonase (Invitrogen). An NRPE1 cDNA fragment corresponding to amino acids 1851-1976 (QS domain) was cloned into the bait vector, pDEST32, and screening against the *A. thaliana* cDNA library was conducted in yeast strain MaV203 (Invitrogen). Screening for interactions using lacZ assays, ura-media, and his- media containing 20mM or 100mM of the HIS3 inhibitor, E-Amino-1, 2, 4 Triazol (3-AT) identified a clone corresponding to RRP6L1 amino acids 421-635. Follow-up Y2H assays, shown in Figure 4, assessed the interaction of the RRP6L1 prey (pDEST22) and NRPE1 bait (pDEST32) corresponding to amino acids 1426-1851 (repeat + DeCL domains), 1426-1651 (repeat domain), and 1851-1976 (QS) domain by detecting growth in the presence of 20mM 3-AT on his- media. Primers utilized for cloning are in Table S5.

### In vitro nuclease activity assays

To test for nuclease activity of heterologously expressed RRP6L1, 50 ng protein was added to a 50 μl reaction containing 1 μl 5 mM nucleic acid substrate and 1X Turbo DNase buffer (Ambion). Reactions were placed at 25°C for 30 minutes and inactivated by cleaning with a Zymogen Oligo Clean and Concentrate Kit according to the manufacturer’s protocol. Samples were visualized on 15% denaturing polyacrylamide gels stained with Sybr Gold (Invitrogen). Negative controls included a no protein control and the addition of an equal volume of nickel column purified protein extract from uninfected insect cells. For positive controls, nucleic acid substrates were also digested with commercially available Turbo DNase (2.5 U/reaction) (Ambion) or RNase A (100 ng/reaction) (Thermo Scientific) at 37°C for 30 minutes. Template sequences are listed in Figure 5B.

### 3’ RACE

RNA was extracted from ~2.5 week old above ground tissues. Since Pol V transcripts lack a polyA tail (Wierzbicki et al. 2008; Wendte and Pikaard 2016), a 3’ poly-A tail was added to 10 gg RNA prior to reverse transcription using commercially available Poly-A polymerase according to the manufacturer’s protocol (NEB). Poly-A tailed RNA was cleaned using the Zymogen RNA clean and concentrate kit. RNA was reverse transcribed using Superscript III (Invitrogen) with a oligo-d(T) primer that added nested PCR primer binding sequences (Table S5). Primary PCR was performed on 1 μl cDNA using Platinum Taq (Invitrogen) according to the manufacturer’s protocol with 3’ RACE Primer 1 and the appropriate gene specific primer (Table S5). Nested PCR was also performed with Platinum Taq on 0.6 μl primary PCR product using 3’ RACE Primer 2 and the appropriate internal gene specific primer (Table S5). PCR conditions for both PCR reactions were: 94°C 2 minutes, 35 cycles of (94°C 30 seconds, 60°C 30 seconds, 72°C 30 seconds). PCR product sizes were verified to be 500 basepairs or less on ethidium bromide-stained agarose gels, pooled, and cleaned using a Qiagen PCR clean-up kit according to the manufacturer’s protocol. For preparation of 3’RACE products for Illumina sequencing, please see supplemental methods.

## Author contributions

J.R.H., J.M.W., and C.S.P. designed the experiments, analyzed the data, and wrote the paper. J.S. conducted *in vitro* transcription assays. O.P. and A.M. conducted immunolocalization assays. J.R.H. and J.M.W. conducted all other experiments.

## Acknowledgements

The authors thank James Ford of the Indiana University Center for Genomics and Bioinformatics for help with library preparation and sequencing, Vibhor Mishra for help with RRP6L1 expression in insect cells, Laurel Bender and Michael Alley of the Indiana University Yeast Two Hybrid Facility, Todd Blevins for sequencing of *nrpe1-11* small RNAs, and Thierry Lagrange for providing seed for ΔSD plants. This work was supported by funds to CSP as an Investigator of the Howard Hughes Medical Institute and Gordon and Betty Moore Foundation and from grant GM077590 from the National Institutes of Health. J.M.W. was supported by NIH training Grant, T32GM007757, and predoctoral fellowship Award F31GM116346. The content of this work is solely the responsibility of the authors and does not necessarily represent the official views of our sponsors. The authors declare no conflicts of interests.

## Supplemental Methods

### Nuclear immunolocalization

For immunolocalization experiments, nuclei from 4-week old plants were fixed in 4% formaldehyde and incubated overnight at 4°C with antibodies recognizing the C-terminal HA epitope of NRPE1 transgene products, as described in (Pontes et al. 2006). Chromatin was counterstained with DAPI.

### Southern blot methylation assays

For the Southern blot analyses described in Figure S2, 250 ng DNA was digested with *Hpa*II or *Hae*III (NEB) and subject to agarose gel electrophoresis and transfer to uncharged nylon membranes. Membranes were probed with a 5S rDNA gene probe generated by random priming of a full length 5S gene PCR product amplified from clone pTC4.2 (Campell et al. 1992).

### RT-PCR

The RT-PCR shown in Figure S2B was conducted using ~1 μg RNA. RNA was treated with RQ1 DNase (Promega) and used to generate random-primed cDNA using degenerate dN6 primers (NEB) and Superscript III Reverse Transcriptase (Invitrogen) according to the manufacturer's instructions. PCR was conducted with GoTaq green (Promega) using primers listed in Table S5. PCR products were analyzed by agarose gel electrophoresis.

### sRNA blot

RNA was isolated from 350 mg inflorescence tissue using a mirVana miRNA isolation kit (Ambion). 9.5 μg RNA was resolved by denaturing polyacrylamide gel electrophoresis on a 20% (w/v) gel. Gels were electroblotted (20 mA/cm2 for 2 hours) to Magnacharge nylon membranes (0.22 μm; Osmonics) using a semidry transfer apparatus. Riboprobes were generated using the mirVana probe construction kit (Ambion) using oligonucleotides specific for a given small RNA and labeling by T7 polymerase transcription in the presence of a-32P CTP. DNA oligonucleotides are listed in Table S5. Blots were hybridized in 50% formamide, 0.25 M Na2HPO4 (pH 7.2), 0.25 M NaCl, 7% SDS at 42°C (14–16 hours) followed by two 15 minute washes at 37°C in 2×SSC, two 15 minute washes at 37°C in 2×SSC, 0.1% SDS, and a 10 minute wash in 0.5×SSC, 1% SDS.

### Heterologous protein expression

*Arabidopsis thaliana* RRP6L1 cDNA, minus the stop codon, was amplified from total RNA using primers described in Table S5. cDNA clones were recombined into the Baculovirus expression system and expressed in Sf9 cells using the Baculo-Direct kit according to the manufacturer’s protocol (Invitrogen). Transformed viruses expressed RRP6L1 modified with a C-terminal 6X His and V5 tag. Protein was purified from P3 virus infected 75 ml Sf9 cultures after incubation for 72 hours. Cells were pelleted by centrifugation at 150 x g for 2 minutes and washed twice with 1X PBS. After the second wash, cells were suspended in binding buffer (0.5 M NaCl, 20 mM Tris-HCl pH 8, 5 mM imidazole) and lysed via sonication with a Biorupter UCD-200 sonicator at 4°C with the settings at Low for 1 minute, and intervals of 10 seconds on, 10 seconds off. Lysate was cleared by centrifugation at 14,000 x g for 20 minutes. Protein was purified from cell extract using Novagen HisBind slurry according to the manufacturer’s protocol. For negative controls, uninfected cells were subject to the same protocol.

### In vitro interaction assays

To test for interaction of insect cell expressed RRP6L1 and the QS subdomain of the NRPE1 CTD, the region encoding amino acids 1851-1976 of the NRPE1 cDNA was amplified and cloned into pDEST15 (Invitrogen) (Table S5). The *β-Glucuronidase* (GUS) reporter gene, also expressed in pDEST15, was used as a negative control. Vectors were expressed *E. coli* BL21-AI One-Shot chemically competent cells grown in LB broth supplemented with 50ug/ml Carbenicillin. For protein expression, 2.5 ml overnight culture was added to 50 ml LB broth (50 μg/ml Carbenicillin), which was incubated at 37°C at 225 rpm until an OD600 was reached (~45 minutes). Expression was induced by the addition of L-arabinose to a concentration of 0.2% and cultures were incubated for a further 3 hours. Cells were then pelleted by centrifugation at 10,000 x g for 10 minutes. Pellets were suspended in 10 ml cold binding buffer (140 mM NaCl, 2.7 mM KCl, 10 mM K_2_HPO_4_, 1.8 mM KH_2_PO_4_, pH 7.4) and lysed via sonication with a Biorupter UDC 200 for 10 minutes on High, 30 seconds on, 30 seconds off. Supernatant was cleared twice by centrifugation for 10 minutes at 10,000 x g and then added to a column containing 300 μl Glutathione Sepharose 4B slurry (Amersham) that had previously been equilibrated with 2 washes of 5 ml binding buffer. Supernatant was incubated with glutathione slurry for 2 hours with end over end rotation at 4°C. Sepharose was washed 2X with 10 ml binding buffer, followed by the addition of 450 ng insect cell expressed RRP6L1 suspended in 10 ml binding buffer. Columns were incubated overnight with end over end rotation at 4°C. Sepharose was then washed 3 times with 10 ml binding buffer. Sepharose or flow-through fractions were suspended in SDS page buffer and western blots were performed as described in the section, *Immunoprecipitation and western blot analysis*, using commercially available anti-GST or anti-V5 antibodies.

### Bisulfite sequencing analyses

Base calling, adapter trimming, and read size selection (>/= 35bp) was performed using bcl2fastq v2.16.0.10. Reads were further quality processed to remove end methylation bias and low quality reads using Cutadapt version 1.9.1 (Martin 2011) with the following commands:

> cutadapt -U 7 -u 2 -q 25

Cleaned reads were mapped to the *Arabidopsis thaliana* TAIR10 genome, followed by the removal of PCR duplicates, and extraction of methylation information for cytosines with a minimum of 5 read coverage using Bismark version 0.16.1 default settings (Krueger and Andrews 2011). The bisulfite conversion rate was calculated based on the number of methylated cytosines divided by total mapped cytosines (converted and un-converted) to the chloroplast genome.

Differently methylated regions (DMRs) were defined using information for cytosine in the CHH context for all genotypes relative to Col-0 using the R package methylKit version 0.9.5 (Akalin et al. 2012). Default methylKit commands were used with the following parameters: The genome was split into 300 base pair sliding windows with a step size of 200 base pairs and a minimum coverage requirement for each window of 10 informative cytosines with at least 5 read coverage each. Significant hypo-DMRs were defined as those regions with a minimum 25% decrease in methylation relative to Col-0 and a q-value of less than or equal to 0.01. To quantify the number of DMRs in each line, overlapping DMRs were merged into a single region. To quantify percent methylation across regions of interest, the methylKit regionCounts function was utilized with the genomic coordinates of interest input as a bed file. In box plots, outliers 1.5 times the interquartile range beyond the upper or lower quartile were omitted. Box plots were generated using the boxplot command in R and heatmaps and clustering analyses were generated using the heatmap2 R package with the default clustering method. Differentially methylated regions defined for each genotype are available in Table S2.

### sRNA sequencing analyses

Raw reads were adapter and quality trimmed and size selected using Cutadapt version 1.9.1 using the following commands:

> cutadapt -a TGGAATTCTCGGGTGCCAAG -q 20 -m 15 -M60

Reads were first filtered of all structural RNAs (tRNAs, rRNAs, snRNAs, and snoRNAs) by mapping to a genome consisting of sequences corresponding to all genomic coordinates identified as structural RNAs on TAIR (www.arabidopsis.org) using bowtie2 default commands. All unmapped reads were saved for further analysis.

Filtered and cleaned reads were mapped to the *Arabidopsis* TAIR10 genome using ShortStack version 3.4 default settings (Johnson et al. 2016). sRNA counts for regions of interest were extracted from bam files using the ShortStack --locifile file function. Counts were normalized as reads per million based on total mapped reads.

### 3’RACE amplicon sequencing

Pooled and cleaned PCR products for each genotype (up to 1.2 μg) were prepared for Illumina sequencing with NEBnext end repair enzyme and Klenow fragment (3’-5’ exo -) from NEB according to the manufacturer’s protocols. After each enzyme treatment, products were cleaned with the Zymogen Clean and Concentrator-5 kit. Adapters (NextFlex PCR free DNA barcodes from BIOO Scientific) were ligated to end repaired, A-tailed PCR products using T4 DNA ligase (NEB) according to the manufacturer’s protocol at 16°C for 2 hours. Libraries were cleaned using 1 volume AMPure XP beads (Beckman Coulter) according to the manufacturer’s instructions and quality checked and quantified with an Agilent Tapestation. 250 basepair single end reads were generated on an Illumina MiSeq by the Indiana University Center for Genomics and Bioinformatics (IU CGB).

Base calling and adapter trimming was performed using bcl2fastq v2.16.0.10 by the IU CGB. Reads were further processed with the following steps to ensure that only reads containing true 3’ end sequences were included in further analyses. Reads were first selected for and trimmed of the 3’ RACE Primer 2 sequence using Cutadapt version 1.9.1 with the following commands:

> cutadapt -g ctactactaggccacgcgtcgactagtac -q 20,20 -m 15 --discard-untrimmed

Reads were then selected for and trimmed of the artificially added poly-A tail with the following:

> cutadapt -g “t(210)” -m 15 --discard-untrimmed

Trimmed reads were then mapped to sequences extracted from the *A. thaliana* TAIR10 genome corresponding to the PCR target regions (consisting of genomic sequences from the following coordinates: Chr4: 2318164-2323164 *(IGN5* top strand); Chr4: 2323282-2328282 (*IGN5* bottom strand); Chr4: 2577970-2582970 (*IGN23*); Chr4: 5459147-5464147 (*IGN25*); Chr2: 15314824-15319824 (*IGN29*); Chr4: 6549057-6554057 (IGN35); Chr1: 13585635-13590635 *(IGN17)* using bowtie2 in local alignment mode (Langmead and Salzberg 2012). SAM output files were converted to bam files, sorted, indexed, and split into individual bam files for each PCR target region using samtools (Li et al. 2009). Bam files were converted to bed files using bedtools (Quinlan and Hall 2010). Bed files were further filtered for high quality mapping scores (MapQ >= 40) and reads that mapped to the appropriate strand, selected for by the GSP. Read coordinates in the bed files were used to calculate the 3’ end lengths relative to the coordinate of the 3’ end of the internal gene-specific primer. Boxplots of relative 3’ end lengths reported in Figures 6 and S5 were generated using the R boxplot function.

---

**Table S1.**
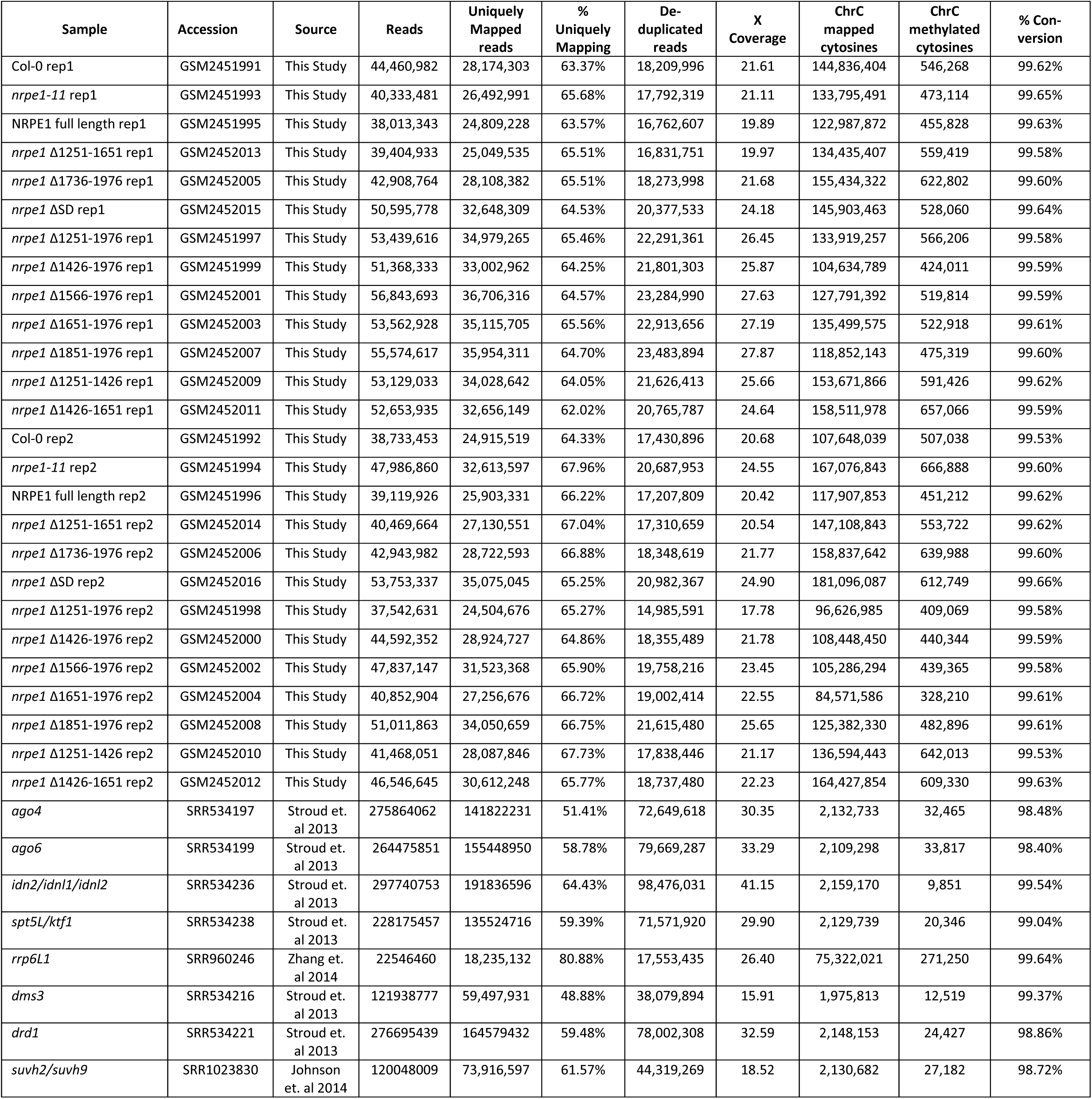
Genome wide bisulfite sequencing stats. Related to Figures 2 and 4.

**Table S2.** Differentially methylated regions. Related to Figures 2, 3, and 4.

**Table S3.** sRNA sequencing data for DMRs. Related to Figure 3F.

**Table S4.**
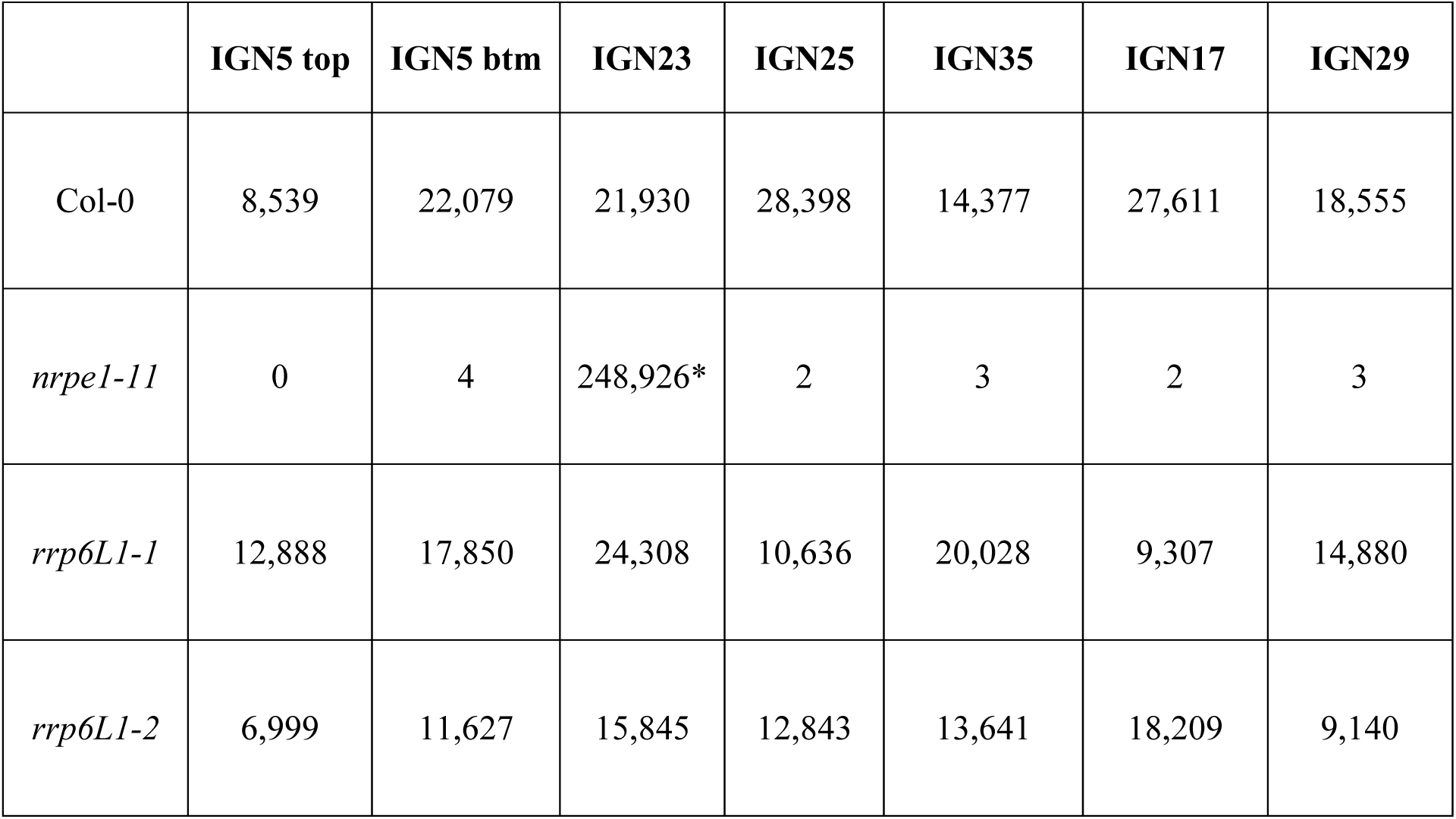
Read counts for 3’ RACE deep sequencing. Related to Figure 6 and S5. *Reads also mapped to a highly similar TE gene, AT3G43686, suggesting a spurious amplification of a derepressed TE in this line. This sequence was filtered from all samples before further analyses

|  | IGN5 top | IGN5 btm | IGN23 | IGN25 | IGN35 | IGN17 | IGN29 |
| --- | --- | --- | --- | --- | --- | --- | --- |
| Col-0 | 8,539 | 22,079 | 21,930 | 28,398 | 14,377 | 27,611 | 18,555 |
| *nrpe1-11* | 0 | 4 | 248,926* | 2 | 3 | 2 | 3 |
| *rrp6L1-1* | 12,888 | 17,850 | 24,308 | 10,636 | 20,028 | 9,307 | 14,880 |
| *rrp6L1-2* | 6,999 | 11,627 | 15,845 | 12,843 | 13,641 | 18,209 | 9,140 |

**Table S5.** Oligonucleotides used in this study. Related to Methods.

**Figure S1.**
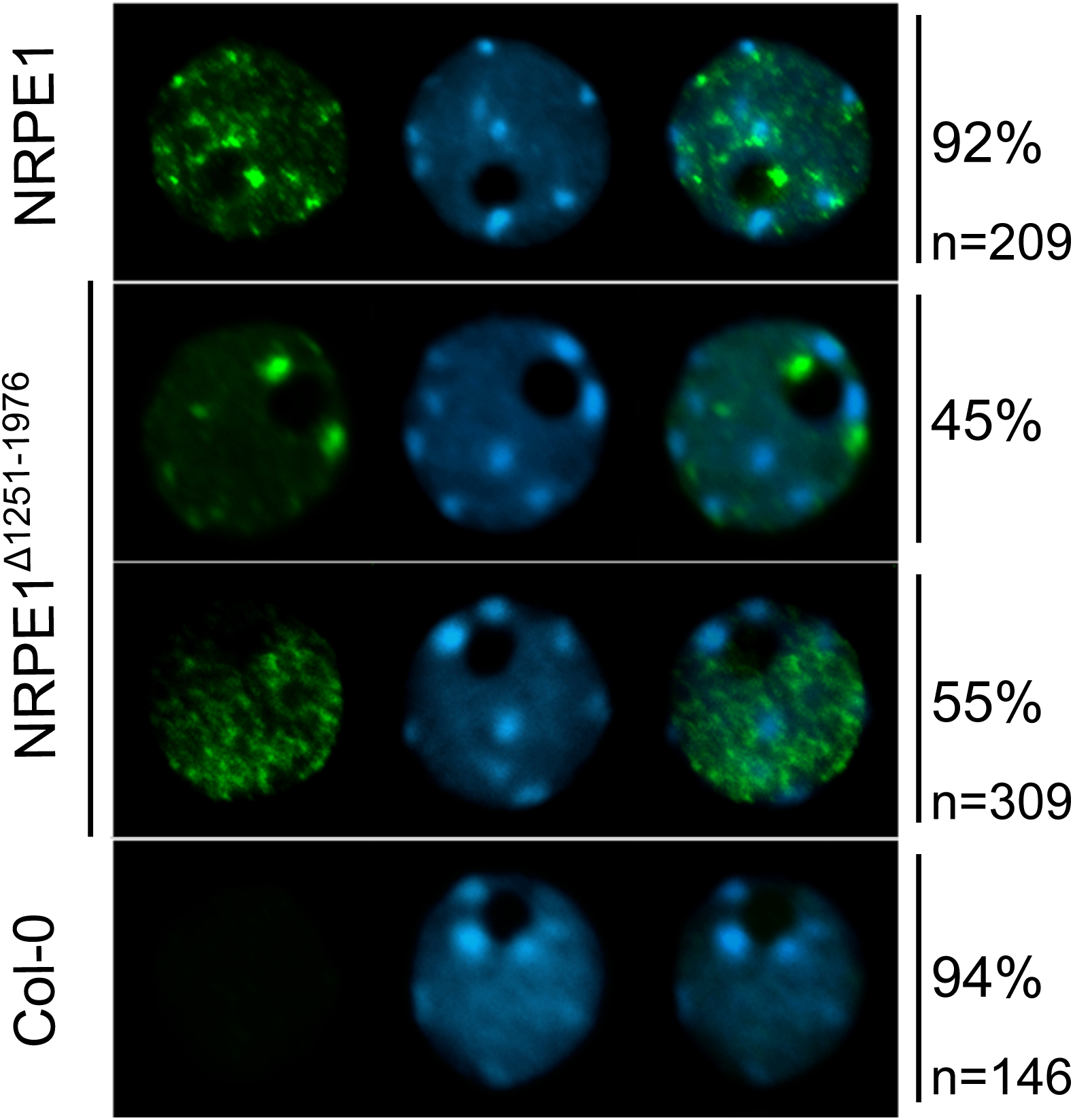
Nuclear immunolocalization of full-length NRPE1 and NRPE1 missing the CTD (Δ1251-1976). Related to Figure 1. Nuclei were fixed using formaldehyde and incubated with anti-HA antibodies recognizing the HA epitope tag at the C-termini of the recombinant proteins. Nuclei of non-transgenic Col-0 serve as a negative control. Nuclei were counterstained with DAPI. Dark, DAPI-negative regions, appearing as black holes, are nucleoli. Bright Pol V foci frequently observed at the edges of the nucleoli (apparent in the top two rows of images) correspond to Cajal bodies. The number of nucleoli examined, and the frequency of localization patterns that resemble the representative images shown, are provided to the right of the images. Note that Pol V assembled using NRPE1 that is missing the CTD displays a nuclear localization pattern only subtly different from that observed for full-length NRPE1. Whereas full-length NRPE1 is present in a larger number of concentrated foci, CTD-deleted NRPE1 tends to be more diffuse, or present in smaller, punctate foci.

**Figure S2.**
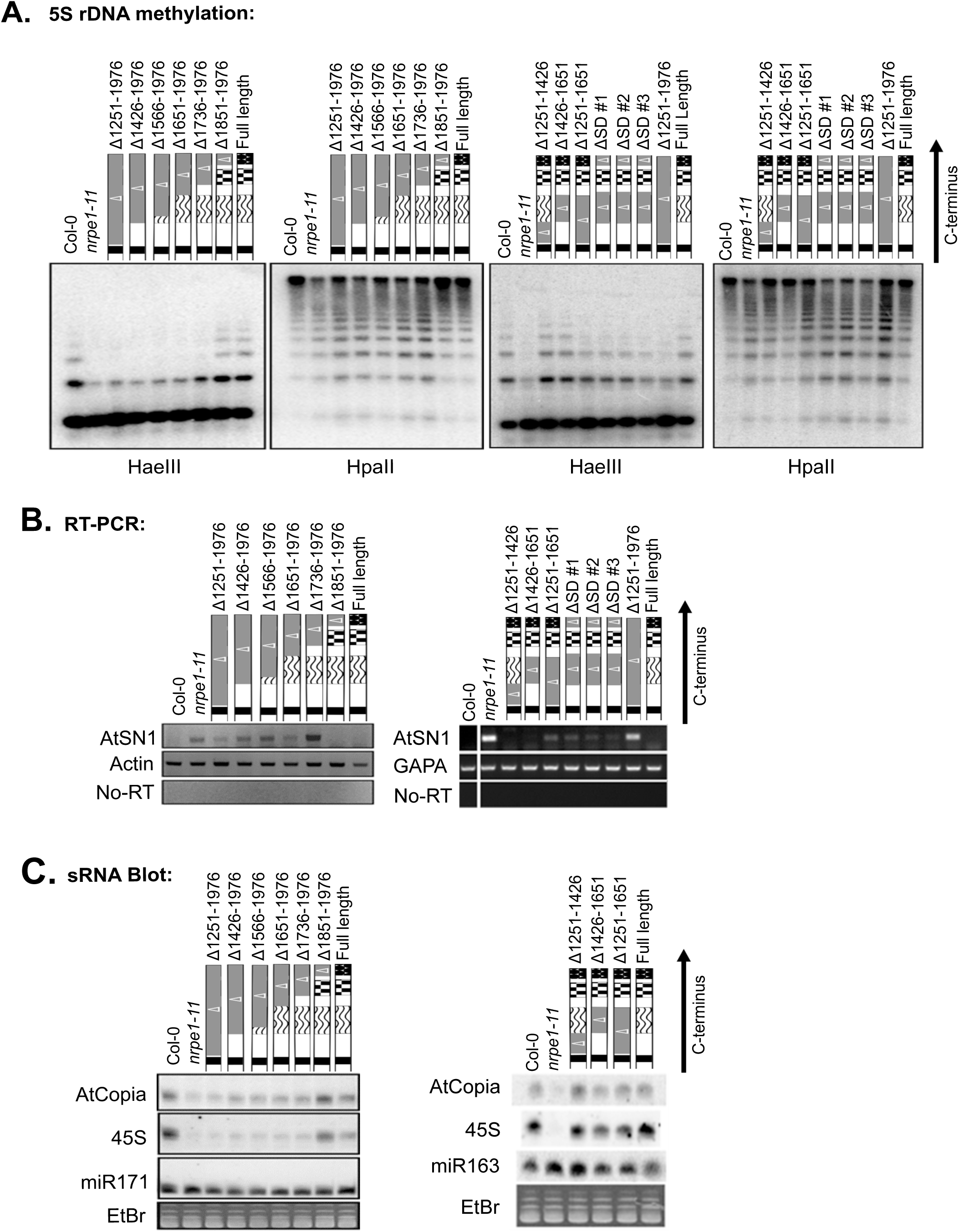
Consequences of NRPE1 CTD deletions on 5S rRNA gene silencing, AtSN1 transposon silencing and siRNA abundance. Related to Figure 2. **A.** Southern blot analysis of 5S rRNA gene repeats after digestion with the methylation sensitive enzymes, *Hae*III or *Hpa*II. Decreased cytosine methylation results in increased digestion, and smaller products. Note the importance of the DeCL domain, and the combinatorial effects of deletions affecting the linker plus 17 aa repeat subdomains or 17aa repeat plus QS subdomains, which have greater effect than deletions of the individual subdomains. **B.** RT-PCR analysis of *AtSN1* retrotransposon transcripts generated when Pol V-dependent silencing is lost. *Actin* and *GAPA* are Pol II transcribed genes included as positive controls. Note that silencing is lost if the DeCL subdomain is deleted. **C.** Northern blot analysis of Pol IV and Pol V-dependent 24nt siRNAs corresponding to *AtCopia* transposons or 45S rRNA gene repeats. *miR171* and *miR163* are 21nt micro RNAs unaffected by the RdDM pathway, included as positive controls. Note that siRNA levels are reduced when the DeCL subdomain is deleted.

**Figure S3.**
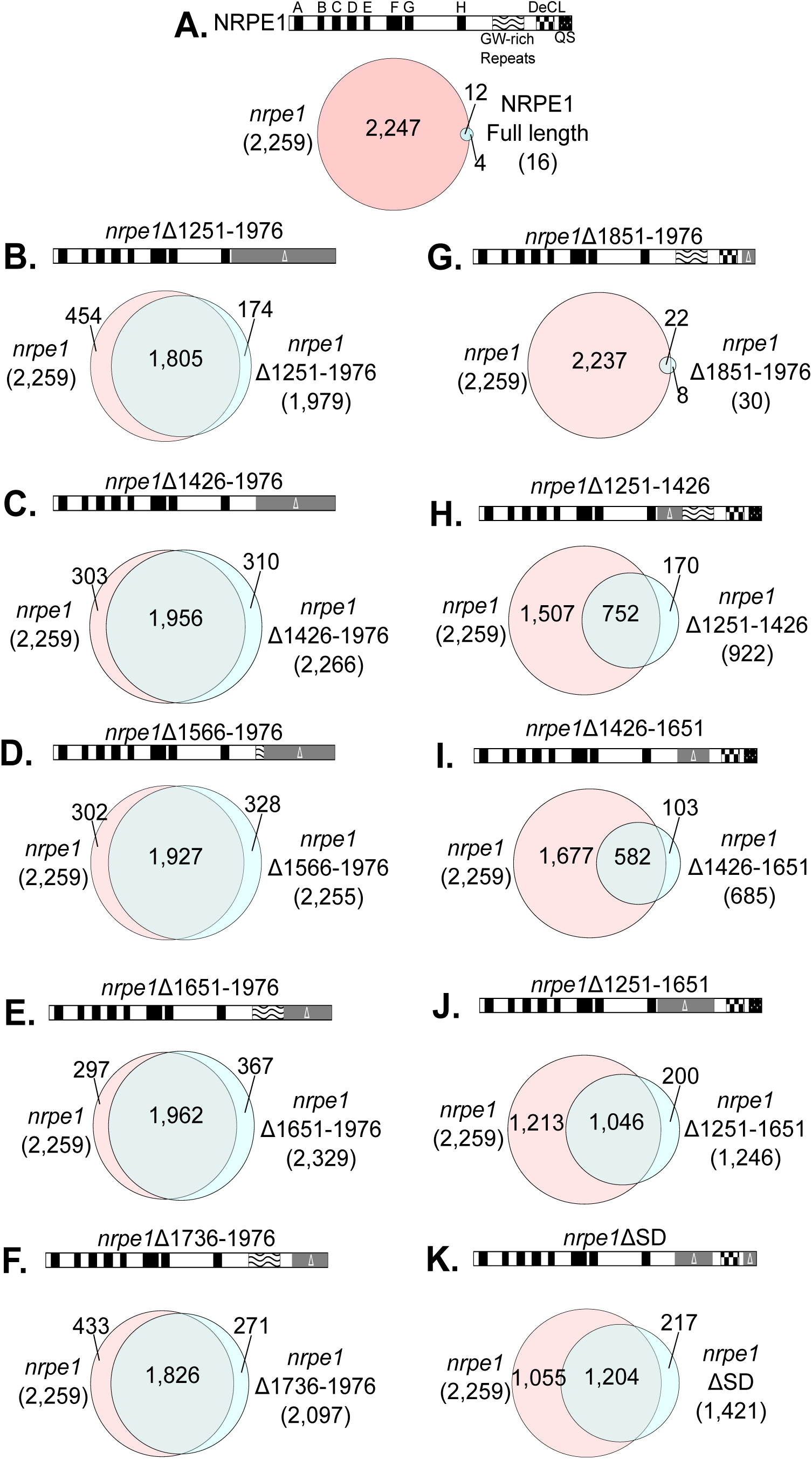
Overlap between CHH DMRs that become hypomethylated in the *nrpe1-11* mutant and DMRs that remain hypomethylated (un-rescued) in each CTD deletion line. Related to Figure 2. All DMRs are defined relative to methylation levels in wild-type Col-0 (see Methods for details).

**Figure S4.**
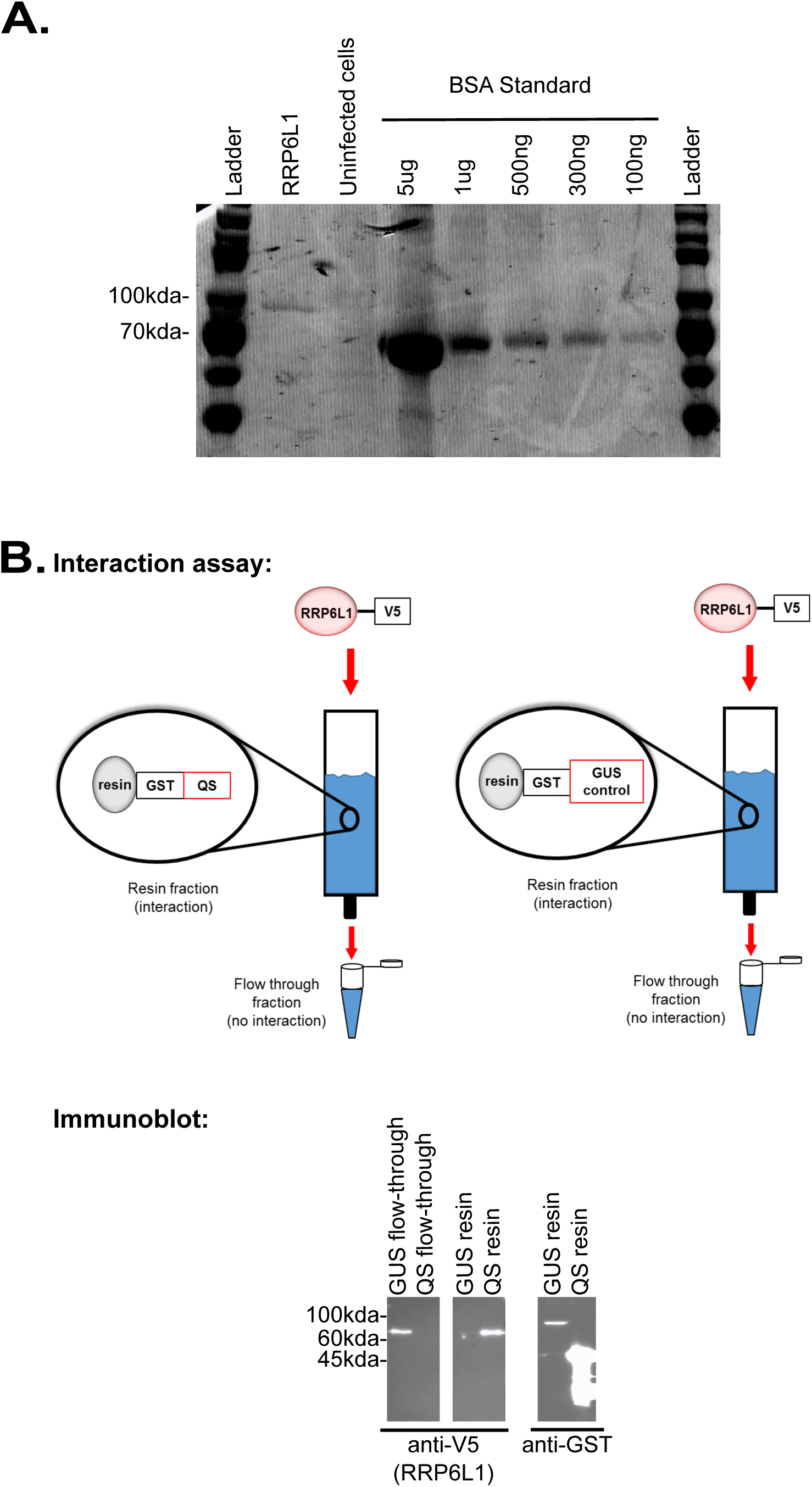
RRP6L1 expressed in insect cells physically interacts with the QS subdomain of NRPE1. Related to Figures 4-5. **A.** Coomassie stained gel showing purified RRP6L1, uninfected cell lysate and a Bovine Serum Albumin (BSA) dilution series. Recombinant RRP6L1 was engineered to have C-terminal His and V5 tags and was expressed in insect cells using a baculovirus vector. The His-tag allowed the protein to be purified using nickel affinity resin. **B.** *In vitro* assay for RRP6L1 interaction with the QS domain (aa1851-1976) of the Pol V CTD. GST-fused to a QS domain polypeptide or a GUS protein control were expressed in bacterial cells and immobilized on glutathione sepharose resin. The resins were then incubated with RRP6L1, and washed extensively to remove unbound protein. RRP6L1 was retained on the resin with immobilized GST-QS peptide, but not the GST-GUS control resin, as shown by SDS-PAGE and immunoblot analyses of resin and flow-through fractions using anti-V5 (recognizing the epitope tag on RRP6L1) or anti-GST antibodies.

**Figure S5.**
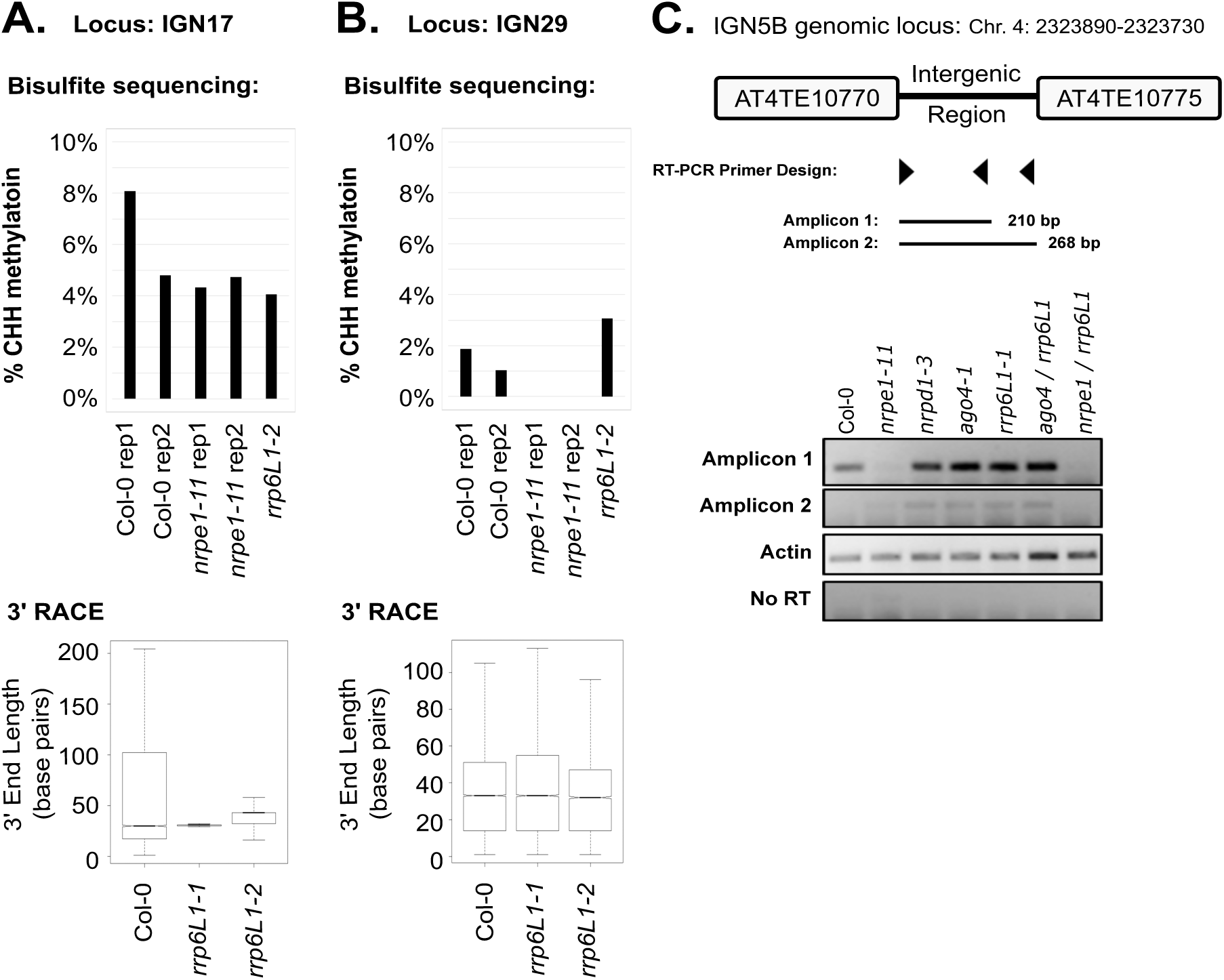
Evidence of RRP6L1 trimming of Pol V transcript 3' ends is not detected at loci where methylation is not dependent on RRP6L1. Related to Figure 6 and Table S4. Bisulfite sequencing and 3’ RACE results for Pol V dependent transcripts from: **A.** *IGN17* (Chr1: 13585614-13585914) and **B.** *IGN29* (Chr2: 15314700-15315000). Histograms show %CHH methylation measured by whole genome bisulfite sequencing. Box plots show the measured 3’ end lengths in basepairs relative to the internal gene specific primer used in the 3’ RACE PCR. **C**. RT-PCR of the Pol V-dependent transcript *IGN5.* Strand-specific RT was performed such that Amplicon 2 detects products with extended 3’ ends.

## References

Ausin I, Greenberg MV, Simanshu DK, Hale CJ, Vashisht AA, Simon SA, Lee TF, Feng S, Espanola SD, Meyers BC et al. 2012. INVOLVED IN DE NOVO 2-containing complex involved in RNA-directed DNA methylation in Arabidopsis. Proc Natl Acad Sci U S A 109: 8374–8381.

Ausin I, Mockler TC, Chory J, Jacobsen SE. 2009. IDN1 and IDN2 are required for de novo DNA methylation in Arabidopsis thaliana. Nat Struct Mol Biol 16: 1325–1327.

Bechtold N, Pelletier G. 1998. In planta Agrobacterium-mediated transformation of adult Arabidopsis thaliana plants by vacuum infiltration. Methods Mol Biol 82: 259–266.

Bellaoui M, Gruissem W. 2004. Altered expression of the Arabidopsis ortholog of DCL affects normal plant development. Planta 219: 819–826.

Bellaoui M, Keddie JS, Gruissem W. 2003. DCL is a plant-specific protein required for plastid ribosomal RNA processing and embryo development. Plant Mol Biol 53: 531–543.

Bies-Etheve N, Pontier D, Lahmy S, Picart C, Vega D, Cooke R, Lagrange T. 2009. RNA-directed DNA methylation requires an AGO4-interacting member of the SPT5 elongation factor family. EMBO Rep 10: 649–654.

Blevins T, Podicheti R, Mishra V, Marasco M, Tang H, Pikaard CS. 2015. Identification of Pol IV and RDR2-dependent precursors of 24 nt siRNAs guiding de novo DNA methylation in Arabidopsis. Elife 4: e09591.

Bohmdorfer G, Rowley MJ, Kucinski J, Zhu Y, Amies I, Wierzbicki AT. 2014. RNA-directed DNA methylation requires stepwise binding of silencing factors to long non-coding RNA. Plant J 79: 181–191.

Chiu J, March PE, Lee R, Tillett D. 2004. Site-directed, Ligase-Independent Mutagenesis (SLIM): a single-tube methodology approaching 100% efficiency in 4 h. Nucleic Acids Res 32: e174.

Clough SJ, Bent AF. 1998. Floral dip: a simplified method for Agrobacterium-mediated transformation of Arabidopsis thaliana. Plant J 16: 735–743.

Earley KW, Haag JR, Pontes O, Opper K, Juehne T, Song K, Pikaard CS. 2006. Gateway-compatible vectors for plant functional genomics and proteomics. Plant J 45: 616–629.

Edgar R, Domrachev M, Lash AE. 2002. Gene Expression Omnibus: NCBI gene expression and hybridization array data repository. Nucleic Acids Res 30: 207–210.

El-Shami M, Pontier D, Lahmy S, Braun L, Picart C, Vega D, Hakimi MA, Jacobsen SE, Cooke R, Lagrange T. 2007. Reiterated WG/GW motifs form functionally and evolutionarily conserved ARGONAUTE-binding platforms in RNAi-related components. Genes Dev 21: 2539–2544.

Fox MJ, Mosley AL. 2016. Rrp6: Integrated roles in nuclear RNA metabolism and transcription termination. Wiley Interdiscip Rev RNA 7: 91–104.

Haag JR, Brower-Toland B, Krieger EK, Sidorenko L, Nicora CD, Norbeck AD, Irsigler A, LaRue H, Brzeski J, McGinnis K et al. 2014. Functional Diversification of Maize RNA Polymerase IV and V Subtypes via Alternative Catalytic Subunits. Cell Rep 9: 378–390.

Haag JR, Pikaard CS. 2011. Multisubunit RNA polymerases IV and V: purveyors of non-coding RNA for plant gene silencing. Nat Rev Mol Cell Biol 12: 483–492.

Haag JR, Ream TS, Marasco M, Nicora CD, Norbeck AD, Pasa-Tolic L, Pikaard CS. 2012. In vitro transcription activities of Pol IV, Pol V, and RDR2 reveal coupling of Pol IV and RDR2 for dsRNA synthesis in plant RNA silencing. Mol Cell 48: 811–818.

He XJ, Hsu YF, Zhu S, Wierzbicki AT, Pontes O, Pikaard CS, Liu HL, Wang CS, Jin H, Zhu JK. 2009. An effector of RNA-directed DNA methylation in arabidopsis is an ARGONAUTE 4- and RNA-binding protein. Cell 137: 498–508.

Huang L, Jones AM, Searle I, Patel K, Vogler H, Hubner NC, Baulcombe DC. 2009. An atypical RNA polymerase involved in RNA silencing shares small subunits with RNA polymerase II. Nat Struct Mol Biol 16: 91–93.

Huang Y, Kendall T, Forsythe ES, Dorantes-Acosta A, Li S, Caballero-Perez J, Chen X, Arteaga-Vazquez M, Beilstein MA, Mosher RA. 2015. Ancient Origin and Recent Innovations of RNA Polymerase IV and V. Mol Biol Evol 32: 1788–1799.

Jing Y, Sun H, Yuan W, Wang Y, Li Q, Liu Y, Li Y, Qian W. 2016. SUVH2 and SUVH9 couple two essential steps for transcriptional gene silencing in Arabidopsis. Mol Plant.

Johnson LM, Du J, Hale CJ, Bischof S, Feng S, Chodavarapu RK, Zhong X, Marson G, Pellegrini M, Segal DJ et al. 2014. SRA- and SET-domain-containing proteins link RNA polymerase V occupancy to DNA methylation. Nature 507: 124–128.

Johnson LM, Law JA, Khattar A, Henderson IR, Jacobsen SE. 2008. SRA-domain proteins required for DRM2-mediated de novo DNA methylation. PLoS genetics 4: e1000280.

Keddie JS, Carroll B, Jones JD, Gruissem W. 1996. The DCL gene of tomato is required for chloroplast development and palisade cell morphogenesis in leaves. Embo J 15: 4208–4217.

Kollen K, Dietz L, Bies-Etheve N, Lagrange T, Grasser M, Grasser KD. 2015. The zinc-finger protein SPT4 interacts with SPT5L/KTF1 and modulates transcriptional silencing in Arabidopsis. FEBS Lett 589: 3254–3257.

Lahmy S, Pontier D, Bies-Etheve N, Laudie M, Feng S, Jobet E, Hale CJ, Cooke R, Hakimi MA, Angelov D et al. 2016. Evidence for ARGONAUTE4-DNA interactions in RNA-directed DNA methylation in plants. Genes Dev 30: 2565–2570.

Lange H, Holec S, Cognat V, Pieuchot L, Le Ret M, Canaday J, Gagliardi D. 2008. Degradation of a polyadenylated rRNA maturation by-product involves one of the three RRP6-like proteins in Arabidopsis thaliana. Mol Cell Biol 28: 3038–3044.

Law JA, Ausin I, Johnson LM, Vashisht AA, Zhu JK, Wohlschlegel JA, Jacobsen SE. 2010. A protein complex required for polymerase V transcripts and RNA- directed DNA methylation in Arabidopsis. Curr Biol 20: 951–956.

Li S, Vandivier LE, Tu B, Gao L, Won SY, Li S, Zheng B, Gregory BD, Chen X. 2015. Detection of Pol IV/RDR2-dependent transcripts at the genomic scale in Arabidopsis reveals features and regulation of siRNA biogenesis. Genome Res 25: 235–245.

Liu ZW, Shao CR, Zhang CJ, Zhou JX, Zhang SW, Li L, Chen S, Huang HW, Cai T, He XJ. 2014. The SET domain proteins SUVH2 and SUVH9 are required for Pol V occupancy at RNA-directed DNA methylation loci. PLoS Genet 10: e1003948.

Luo J, Hall BD. 2007. A multistep process gave rise to RNA polymerase IV of land plants. J Mol Evol 64: 101–112.

Matzke MA, Mosher RA. 2014. RNA-directed DNA methylation: an epigenetic pathway of increasing complexity. Nat Rev Genet 15: 394–408.

McCue AD, Panda K, Nuthikattu S, Choudury SG, Thomas EN, Slotkin RK. 2015. ARGONAUTE 6 bridges transposable element mRNA-derived siRNAs to the establishment of DNA methylation. EMBO J 34: 20–35.

Mosher RA, Schwach F, Studholme D, Baulcombe DC. 2008. PolIVb influences RNA-directed DNA methylation independently of its role in siRNA biogenesis. Proc Natl Acad Sci U S A 105: 3145–3150.

Murray MG, Thompson WF. 1980. Rapid isolation of high molecular weight plant DNA. Nucleic Acids Res 8: 4321–4325.

Nuthikattu S, McCue AD, Panda K, Fultz D, DeFraia C, Thomas EN, Slotkin RK. 2013. The initiation of epigenetic silencing of active transposable elements is triggered by RDR6 and 21-22 nucleotide small interfering RNAs. Plant Physiol 162: 116–131.

Onodera Y, Haag JR, Ream T, Costa Nunes P, Pontes O, Pikaard CS. 2005. Plant nuclear RNA polymerase IV mediates siRNA and DNA methylation-dependent heterochromatin formation. Cell 120: 613–622.

Pontes O, Costa-Nunes P, Vithayathil P, Pikaard CS. 2009. RNA polymerase V functions in Arabidopsis interphase heterochromatin organization independently of the 24-nt siRNA-directed DNA methylation pathway. Mol Plant 2: 700–710.

Pontes O, Li CF, Costa Nunes P, Haag J, Ream T, Vitins A, Jacobsen SE, Pikaard CS. 2006. The Arabidopsis chromatin-modifying nuclear siRNA pathway involves a nucleolar RNA processing center. Cell 126: 79–92.

Pontier D, Picart C, Roudier F, Garcia D, Lahmy S, Azevedo J, Alart E, Laudie M, Karlowski WM, Cooke R et al. 2012. NERD, a plant-specific GW protein, defines an additional RNAi-dependent chromatin-based pathway in Arabidopsis. Mol Cell 48: 121–132.

Pontier D, Yahubyan G, Vega D, Bulski A, Saez-Vasquez J, Hakimi MA, Lerbs-Mache S, Colot V, Lagrange T. 2005. Reinforcement of silencing at transposons and highly repeated sequences requires the concerted action of two distinct RNA polymerases IV in Arabidopsis. Genes Dev 19: 2030–2040.

Qi Y, He X, Wang XJ, Kohany O, Jurka J, Hannon GJ. 2006. Distinct catalytic and non-catalytic roles of ARGONAUTE4 in RNA-directed DNA methylation. Nature 443: 1008–1012.

Ream T, Haag J, Pikaard CS. 2013. Plant multisubunit RNA polymerases IV and V. in Nucleic Acid Polymerases (eds. K Murakami, M Trakselis), pp. 289–308. Springer-Verlag, Berlin Heidelberg.

Ream TS, Haag JR, Wierzbicki AT, Nicora CD, Norbeck AD, Zhu JK, Hagen G, Guilfoyle TJ, Pasa-Tolic L, Pikaard CS. 2009. Subunit compositions of the RNA-silencing enzymes Pol IV and Pol V reveal their origins as specialized forms of RNA polymerase II. Mol Cell 33: 192–203.

Rowley MJ, Avrutsky MI, Sifuentes CJ, Pereira L, Wierzbicki AT. 2011. Independent chromatin binding of ARGONAUTE4 and SPT5L/KTF1 mediates transcriptional gene silencing. PLoS Genet 7: e1002120.

Rowley MJ, Bohmdorfer G, Wierzbicki AT. 2013. Analysis of long non-coding RNAs produced by a specialized RNA polymerase in Arabidopsis thaliana. Methods 63: 160–169.

Stroud H, Greenberg MV, Feng S, Bernatavichute YV, Jacobsen SE. 2013. Comprehensive analysis of silencing mutants reveals complex regulation of the Arabidopsis methylome. Cell 152: 352–364.

Trujillo JT, Beilstein MA, Mosher RA. 2016. The Argonaute-binding platform of NRPE1 evolves through modulation of intrinsically disordered repeats. New Phytol 212: 1094–1105.

Tucker SL, Reece J, Ream TS, Pikaard CS. 2010. Evolutionary history of plant multisubunit RNA polymerases IV and V: subunit origins via genome-wide and segmental gene duplications, retrotransposition, and lineage-specific subfunctionalization. Cold Spring Harb Symp Quant Biol 75: 285–297.

Wang Y, Ma H. 2015. Step-wise and lineage-specific diversification of plant RNA polymerase genes and origin of the largest plant-specific subunits. New Phytol 207: 1198–1212.

Wendte JM, Pikaard CS. 2016. The RNAs of RNA-directed DNA methylation. Biochim Biophys Acta.

Wierzbicki AT, Cocklin R, Mayampurath A, Lister R, Rowley MJ, Gregory BD, Ecker JR, Tang H, Pikaard CS. 2012. Spatial and functional relationships among Pol V-associated loci, Pol IV-dependent siRNAs, and cytosine methylation in the Arabidopsis epigenome. Genes Dev 26: 1825–1836.

Wierzbicki AT, Haag JR, Pikaard CS. 2008. Noncoding transcription by RNA polymerase Pol IVb/Pol V mediates transcriptional silencing of overlapping and adjacent genes. Cell 135: 635–648.

Wierzbicki AT, Ream TS, Haag JR, Pikaard CS. 2009. RNA polymerase V transcription guides ARGONAUTE4 to chromatin. Nature genetics 41: 630–634.

Zaborowska J, Egloff S, Murphy S. 2016. The pol II CTD: new twists in the tail. Nat Struct Mol Biol 23: 771–777.

Zhai J, Bischof S, Wang H, Feng S, Lee TF, Teng C, Chen X, Park SY, Liu L, Gallego-Bartolome J et al. 2015. A One Precursor One siRNA Model for Pol IV-Dependent siRNA Biogenesis. Cell 163: 445–455.

Zhang CJ, Ning YQ, Zhang SW, Chen Q, Shao CR, Guo YW, Zhou JX, Li L, Chen S, He XJ. 2012. IDN2 and its paralogs form a complex required for RNA-directed DNA methylation. PLoS Genet 8:e1002693.

Zhang H, Tang K, Qian W, Duan CG, Wang B, Zhang H, Wang P, Zhu X, Lang Z, Yang Y et al. 2014. An Rrp6-like protein positively regulates noncoding RNA levels and DNA methylation in Arabidopsis. Molecular cell 54: 418–430.

Zhang X, Henderson IR, Lu C, Green PJ, Jacobsen SE. 2007. Role of RNA polymerase IV in plant small RNA metabolism. Proc Natl Acad Sci U S A 104: 4536–4541.

Zhong X, Hale CJ, Law JA, Johnson LM, Feng S, Tu A, Jacobsen SE. 2012. DDR complex facilitates global association of RNA polymerase V to promoters and evolutionarily young transposons. Nat Struct Mol Biol 19: 870–875.

Zhou M, Law JA. 2015. RNA Pol IV and V in gene silencing: Rebel polymerases evolving away from Pol II's rules. Curr Opin Plant Biol 27: 154–164.

## References

Akalin A, Kormaksson M, Li S, Garrett-Bakelman FE, Figueroa ME, Melnick A, Mason CE. 2012. methylKit: a comprehensive R package for the analysis of genome-wide DNA methylation profiles. Genome Biol 13: R87.

Campell BR, Song Y, Posch TE, Cullis CA, Town CD. 1992. Sequence and organization of 5S ribosomal RNA-encoding genes of Arabidopsis thaliana. Gene 112: 225–228.

Johnson NR, Yeoh JM, Coruh C, Axtell MJ. 2016. Improved Placement of Multi-mapping Small RNAs. G3 (Bethesda) 6: 2103–2111.

Krueger F, Andrews SR. 2011. Bismark: a flexible aligner and methylation caller for Bisulfite-Seq applications. Bioinformatics 27: 1571–1572.

Langmead B, Salzberg SL. 2012. Fast gapped-read alignment with Bowtie 2. Nat Methods 9: 357–359.

Li H, Handsaker B, Wysoker A, Fennell T, Ruan J, Homer N, Marth G, Abecasis G, Durbin R, Genome Project Data Processing S. 2009. The Sequence Alignment/Map format and SAMtools. Bioinformatics 25: 2078–2079.

Martin M. 2011. Cutadapt removes adapter sequences from high-throughput sequencing reads. EMBnetjournal 17: 10–12.

Quinlan AR, Hall IM. 2010. BEDTools: a flexible suite of utilities for comparing genomic features. Bioinformatics 26: 841–842.

